# General solution to biological signalling games: costly signalling and beyond

**DOI:** 10.1101/2022.05.10.491297

**Authors:** Szabolcs Számadó, István Zachar, Dániel Czégel, Dustin J. Penn

## Abstract

Explaining signal reliability poses a central problem in animal communication. According to Zahavi’s Handicap Principle (HP), signals are honest only when they are costly at the evolutionary equilibrium – hence the term ‘handicap’; otherwise, deception evolves, and communication breaks down. The HP has no theoretical or empirical support, despite claims to the contrary, and yet this idea remains immensely popular. Theoretical evaluations of the HP are difficult, however, because finding the equilibrium cost function in signalling games is notoriously complicated. Here we show how cost functions can be calculated for any arbitrary pairwise asymmetric signalling game at the evolutionary equilibrium. We clarify the relationship between signalling costs at equilibrium and the conditions for reliable signalling. We show that these two terms are independent and the costs of signalling at honest equilibrium have no effect on the stability of communication. We show that honest signals can take any cost value, even negative, being beneficial for the signaller independently of the receiver’s response at equilibrium, without requiring further constraints. Our results are general, and apply to seminal signalling models, including Grafen’s model of sexual selection and Godfray’s model of parent-offspring communication. Our results refute the claim that signals must be costly at the evolutionary equilibrium to be reliable, as predicted by the HP and so-called “costly signalling” theory. The handicap paradigm can thus be fully rejected. We provide testable predictions to help advance the field and establish a better explanation for honest signals.

## Introduction

Explaining the evolution of honest signalling has been a long-standing problem in research on animal^1^ and human communication^2^. Zahavi’s Handicap Principle^3,4^ is the leading theoretical paradigm for honest signalling, and it predicts that signals must be costly at the evolutionary equilibrium – hence the name ‘handicap’ – in order to be honest. This idea is often claimed to provide a general principle to explain why signals are honest, although some have questioned its generality^5^. There is no consensus for how to define, model or test the Handicap Principle, or related ideas known as “costly signalling theory” because ‘handicaps’ and signalling costs have never been clearly defined, nor shown how they enforce honesty by their proponents^5–8^.

Mathematical signalling games have greatly improved our understanding of honest signalling^1,9,10^. They have helped to clarify the logic of honest signalling in conspecific interactions, including aggression^11^, mate choice^12^, parent-offspring conflict^13–15^, and interspecific interactions, such as plant-herbivore^16^, plant-pollinator^17^, aposematic displays^18^, and predator-prey^19^ relations. They originated with economic signalling games^20^ and have been used to analyse the stability of honest signals in a variety of human social interactions^21^. However, identifying the costs of signalling at the honest evolutionary equilibrium (equilibrium cost function) in such models is not trivial when the signallers’ quality can vary continuously^12,15^.

The most influential model of honest signalling is Grafen’s so-called “strategic handicap” model critically assumes that signallers differ in their quality but face marginal, differential costs for producing a signal, so that there is a fitness cost for dishonest signalling. widely misinterpreted model, because it is not a handicap model^5^. Unlike the Handicap Principle, signals in Grafen’s model are efficient rather than wasteful, and honesty does not require high absolute signalling costs. Moreover, honest signals are selectively favoured in the model despite of their costs; not because they are costly. This model has nevertheless provided an important step towards analysing fitness trade-offs for honest signalling, but the steps used by Grafen to obtain the equilibrium cost function are difficult to replicate, hence his mathematics have been described as ‘arcane’^7^. It is also incomplete: finding an equilibrium cost function in general requires solving a double optimization problem^13,22^ – one for the receiver and another for the signaller, which depends on the solution to the former – this complexity of often prohibiting and thus sometimes circumvented by ignoring receiver’s optimization problem (see Appendix 5-7 for a more detailed discussion). This issue cannot be resolved until the optimization problem of the signaller and the receiver are both evaluated.

This double optimization problem has an infinite number of possible solutions^22,23^, and no general solution has ever been provided in analytical form. The lack of a clear methodology for deriving solutions to address this double optimization problem contributes to widespread misinterpretations of the Handicap Principle and Grafen’s model (see^5^), leaving critiques difficult to understand^6,7^. It has been known for 20 years that such signalling games have an infinite number of honest equilibria^22,23^, and yet the nature and implications of these equilibria have remained unexplored due to the complexity of this challenging problem. Consequently, the conditions for honest communication in signalling games is still unclear and controversial, and the field has stagnated due to being entwined in the erroneous and confusing handicap paradigm^5^.

Here, we provide a novel, general approach for determining stable equilibria in continuous signalling games, and for calculating equilibrium signal cost functions – as a continuation of previous theoretical developments^24^. We examine signalling models with *additive fitness functions* (when signal costs and benefits are measured in the same currency, such as fitness), and also *multiplicative fitness functions* (such as when signals have survival costs that influence the reproductive benefits)^25^. First, we describe an asymmetric signalling model of animal communication, general enough to apply to any context in which certain conditions are met. Then we provide solutions for games with additive or multiplicative fitness functions. We provide a formal proof of the conditions for stability being independent of equilibrium signal cost. Our general formula specifies the full, infinite set of trade-off solutions. Furthermore, we show that an infinite number of cost-free and negative cost equilibria exist in these models. The discovery of these previously unknown and evolutionarily stable equilibria shows how new approaches and interpretations can be used to investigate signalling games in general.

Our approach does not require prior knowledge or assumptions about the shape of the potential solution, and hence it is applicable to any model. We apply our method to calculate equilibria in classic signalling models, including Grafen’s model for sexual signalsGodfray’s signal-of-need model^15^ for parent-offspring signalling games, and for the signalling model of Bergstrom et al.^22^. We explain our results regarding cost-free and beneficial (negative-cost) honest signals at equilibrium and provide testable predictions that could support (or refute) our results and their generality. Finally, we explain the concept of beneficial signals and how they can be integrated into honest signalling theory.

## Background

Signalling games are mathematical models used to analyse how individuals (*signallers)* attempt to influence the decisions of others (*receivers)* by producing signals (action at a distance, see Fig. 1). ‘Signals’ are often defined as traits that provide information about some aspect of a signaller that is not directly observable, such as size or sex^20^. Signalling games are usually described as conflicts over a resource, because some of the first models were contests over food and territories. Indeed, from an evolutionary perspective, a receiver’s body and behaviour can be viewed as scarce resources over which signallers compete to exploit for their own benefit^26^. We adopt an evolutionary conflict perspective here, though this does not imply that altruistic actions cannot be explained by natural selection and provide fitness benefits. Signalling games can be symmetric or asymmetric concerning information, resources, and options (strategies) available to the players. In symmetric games, players have the same information sets, resource availability and strategies at the beginning of the game, whereas in asymmetric games, players do not share the same information, resources, or strategic options.

**Fig. 1.**
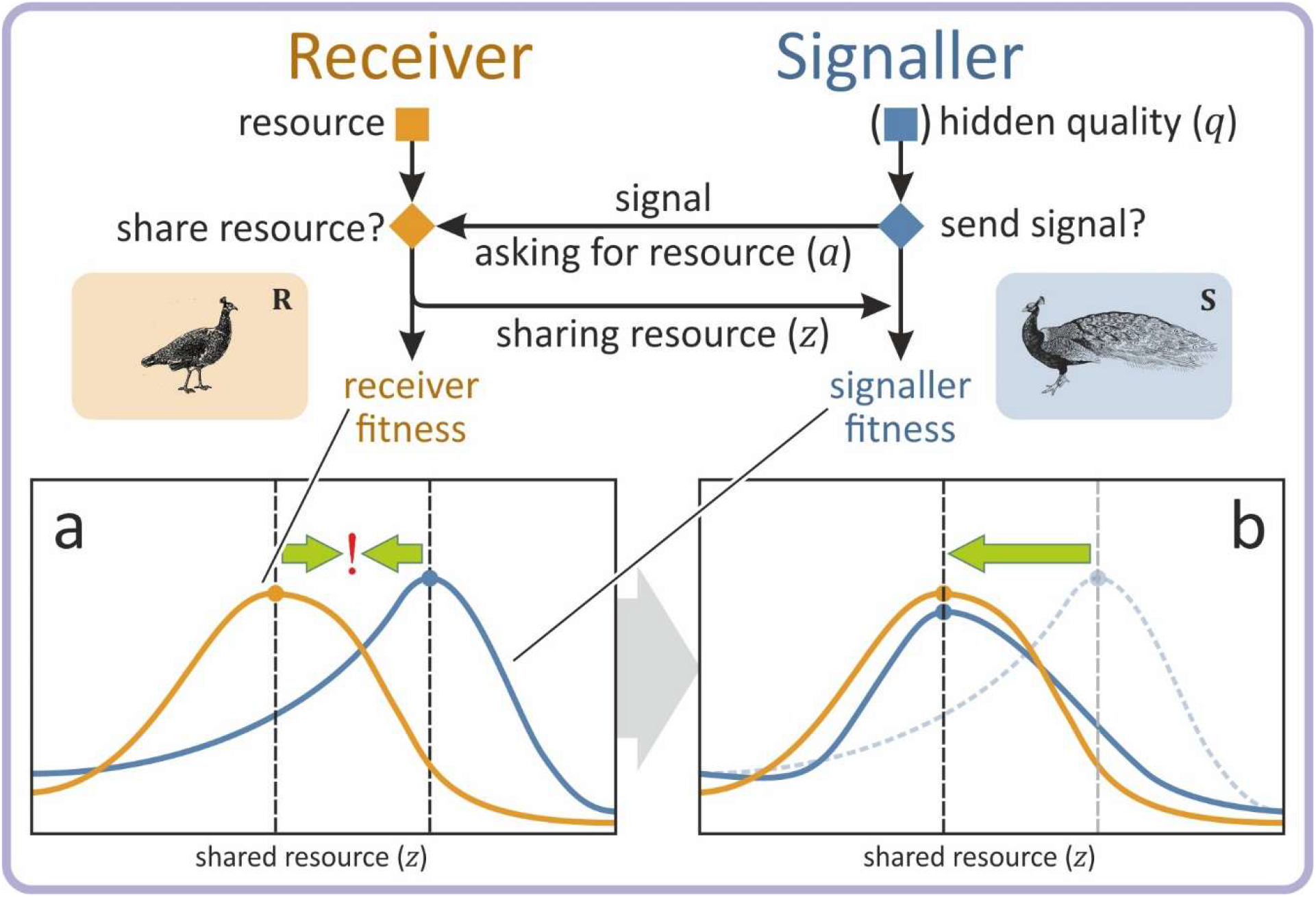
Female mate choice: an asymmetric signalling game. Courtship and mating behaviour is a sexual interaction in which a male signaller (S) aims to persuade a choosey female receiver (R) to mate and allow him to fertilise her eggs; a scarce resource (e.g.^12^). Males vary in their quality and R will mate with a S depending upon his quality. S knows his own quality, whereas R cannot evaluate his quality directly; she must decide based solely on attributes of S’s signal (i.e., there is an information asymmetry). Axes *x* and *y* in the inset images represent the amount of shared resource *z* and fitness w outcomes respectively. Inset **a**: For a given signaller quality and no signalling trade-off, the fitness curves of S (blue) and R (yellow) have their optima (blue and yellow points) at different amounts of shared resource (dashed lines), resulting in a conflict of interest. Inset **b**: Signal trade-offs modify the signaller’s fitness curve (blue) such that its optimum is at the same amount of resource that the receiver is willing to share, reducing or eliminating the conflict in the interaction. Costly signalling theory predicts that this trade-off function must be positive at the equilibrium for signalling to be honest^12,15^.

Games can be symmetrical or asymmetrical in many respects, and the consequences of information asymmetries have long attracted the interest of biologists^27^ and economists^20,28^. Asymmetrical signalling games are often used to understand how individuals resolve *conflicts*, and biologists have used them to model a wide variety of interactions and types of conflicts (e.g., genomic, sexual, parent-offspring, and other intra- and inter-familial conflicts). In these models, signallers benefit by persuading a receiver to take some action, which can include mating^12^, feeding^15^, other forms of parental investment^29^, committing suicide^30,31^, or performing other actions that may or may not be in the receiver’s interest. Asymmetrical signalling games have been used to model intra-genomic conflicts and molecular signals between cells within the body^32,33^. They have also been used to model a variety of interspecific interactions, including predator-prey^19,34^, host-parasite^35–37^, plant-herbivore^16^, plant-pollinator^17^ and aposematic displays^18^. They are also used to model and combat the spread of misinformation in human societies, which arguably is a crucial issue^38–40^.

Here we focus on games with asymmetries in access to both information and resources. In asymmetric games, receivers possess a resource, and can decide whether to share it with signallers or not. For example, young chicks attempt to persuade their parents to feed them by producing begging calls^15^. In discrete models, receivers can either give away the entire resource or keep it for themselves^9,16,19,41–43^, whereas in continuous models, receivers can share some **portion of the resource** (*z*)^12,13,15,22–25,44^. Receivers are assumed to share the resource in a way that maximizes (inclusive) **receiver fitness** (*w*_**R**_), but the potential benefits depend upon obtaining reliable information from signallers about what they offer in exchange. The problem is that receivers often have incomplete information about signallers or what they have to offer (information asymmetry). In the case of mate choice, females assess the potential benefits of mating with males that differ in social status, health, resources, or other aspects of **quality** (*q*); however, male quality cannot be directly assessed by the receiver, otherwise there is no need for signals. The signaller can influence the receiver’s decision by its signal, which may or may not reliably reveal the quality *q* of the signaller, to **ask for the resource amount** (*a*) that should maximize **signaller fitness** (*w*_**S**_) (see Fig. 1). A signal is honest if it enables receivers to obtain reliable information about the signaller’s quality, allowing the receiver to make adaptive decisions. Alternatively, the signal can be useless or deceptive, so that signallers manipulate the receivers to share more than the amount *z* that is in their adaptive interest. Honest signalling models investigate the conditions under which signals are reliable indicators of quality *q* (for details, see Methods and Appendix 1-3).

Theoretical models have previously shown that honest signals are evolutionarily stable at an *honest equilibrium* if the following conditions are met^13,22^: (*i*) the signal reveals the signaller’s actual quality *q* (signals are honest), so that the receiver can respond adaptively; or (*ii*) the signaller only asks for the amount *a* of a resource that receivers benefit by sharing (shared interest), so that the conflict between the receiver and signaller is removed at the honest equilibrium (*a* = *z*). The mathematical formulation of these conditions is detailed in Methods.

The standard theoretical approach used to resolve conflicts of interest in asymmetric signalling games is to introduce a *cost function* that transforms the signaller’s fitness function *w*_**S**_, so that its optimal amount of resource *a* acquired by the signaller corresponds to the optimal amount of resource that the receiver shares *z* (see Fig. 1). We refer to this cost function as *T* for *trade-off function* because the term ‘cost function’ is unnecessarily restrictive to the positive domain (cost value is positive), and does not represent the full set of solutions, as we will demonstrate below.

Accordingly, the signaller’s fitness *w*_**S**_ consists of a benefit *B* and a trade-off function *T*; and without trade-offs, *w*_**S**_ = *B*. In additive models (e.g.^15^), *B* and *T* are summed, whereas in multiplicative models (e.g.^12^), they are multiplied to yield the fitness *w*_**S**_.

In this paper, we construct the most general class of trade-off functions that obey the conditions of honest signalling for both additive and multiplicative fitness functions (see Methods and Appendices 2-3). Lastly, we apply our method to well-known models of honest signalling (Appendix 4), demonstrating its general applicability.

## Results

Since it is the fitness *w*_**S**_ that must meet the conditions of stability and honesty in an honest equilibrium, and not the benefit *B*, we first show that a signal trade-off function *T* can always be found for any *B* that ensures that *w*_**S**_ meets these conditions. In order to decompose the signaller’s fitness function into terms that are in one-to-one correspondence with the conditions of honest signalling, we expand the signaller’s fitness function to its Taylor-series around (honest) signalling equilibrium. This representation allows us the derive the exact and most general implications of the conditions of honest signalling term by term. As we show below, the conditions of honest signalling constrain the first order and second order terms, while the rest can be chosen arbitrarily. When terms are again summed up, the resulting *w*_**S**_ represents all honest solutions of signalling. **Fig. 2** illustrates the process for additive and multiplicative models, while Fig. 3 provides a visual guide for the method of constructing *T* for the additive case (see Methods, Appendix 2 and Fig. S1).

**Fig. 2.**
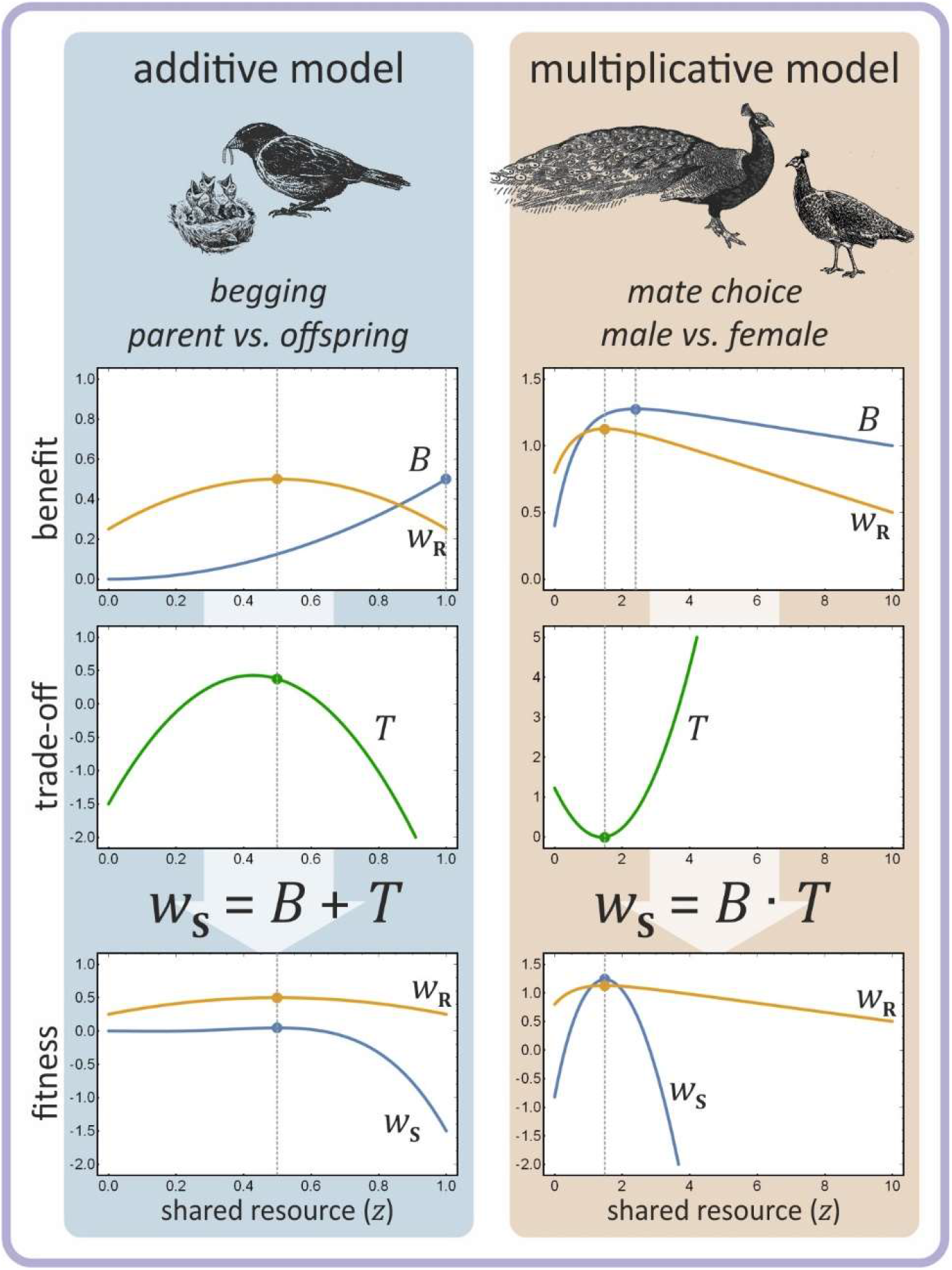
Cost/benefit trade-off functions for two traditional signalling games. Left: An additive offspring begging game. Right: A multiplicative mate choice signalling game. Dashed vertical lines indicate the receiver’s and the signaller’s optimal amount of resource, given signaller’s quality. Trade-off function *T*, when added to or multiplied by *B*, transforms the signaller’s benefit *B* to its actual fitness *w*_**S**_ such that its optimum amount of requested resource *a* coincides with the amount *z* shared by the receiver.

**Fig. 3.**
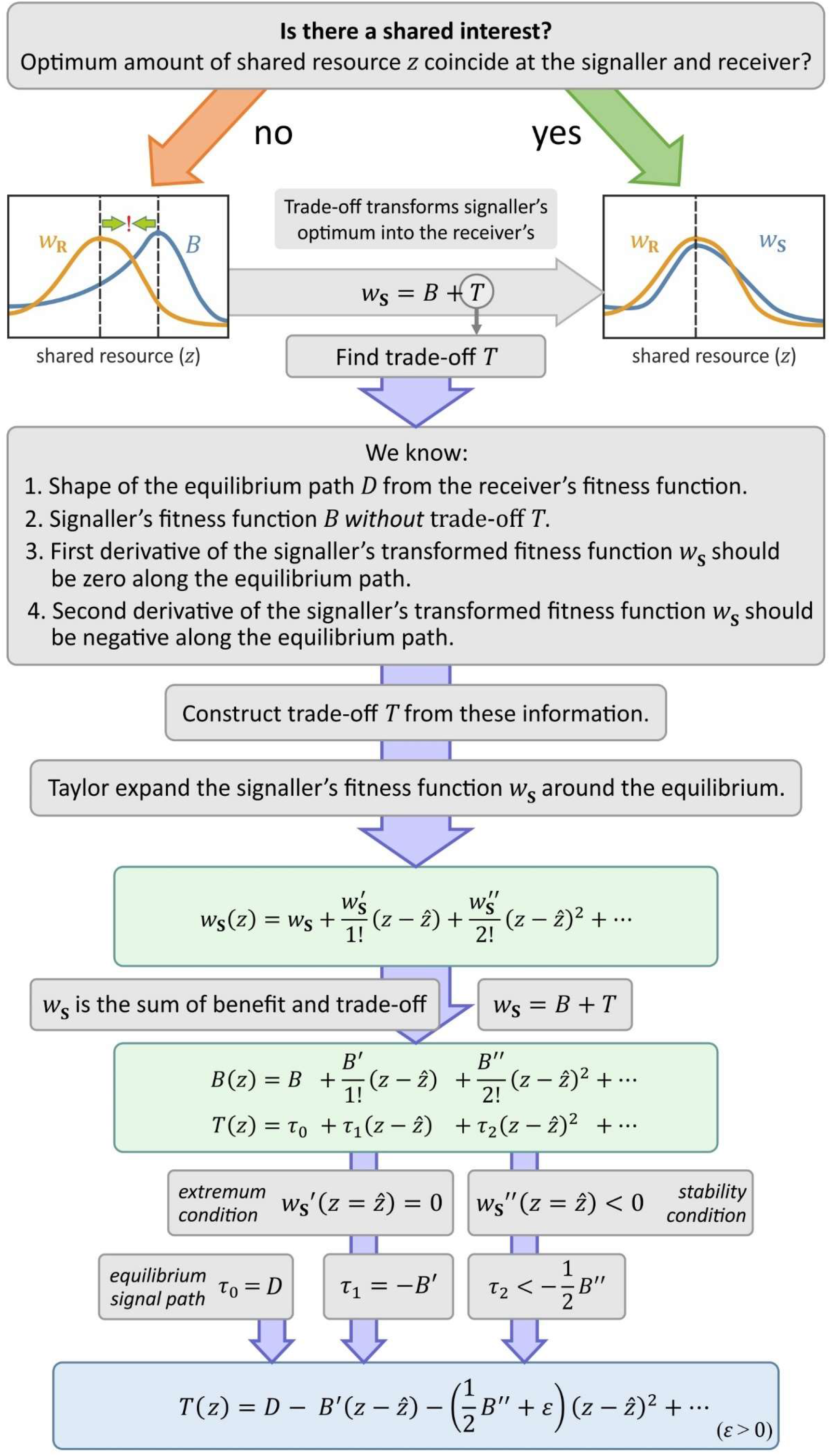
Method of reverse-engineering the general trade-off function *T*. The method transforms any (at least twice) differentiable signaller benefit function *B* to the fitness function *w*_**S**_ that has the same optimal amount of shared resource as the receiver’s fitness function *w*_R_. For sake of simplicity, function arguments are omitted at the right side of equations. *D* and higher order Taylor coefficients *τ*_3_, *τ*_4_, … can be freely chosen.

### Additive fitness

The general form of any (at least twice differentiable) trade-off function for additive fitness *T*_*A*_ using Taylor-series expansion around the equilibrium where 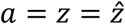 is (see Appendix 2 and Fig. S1 for details):

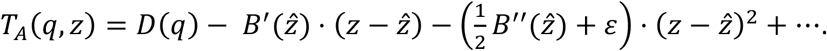

The zeroth order Taylor coefficient *D*(*q*) is the *equilibrium signal trade-off function* of offspring quality *q*. Traditionally, this coefficient specifies the cost that signallers pay at the equilibrium, independently of the conditions of honest signalling^15^. **Fig. 4** shows a set of example costly and cost-free equilibrium cost functions.

**Fig. 4.**
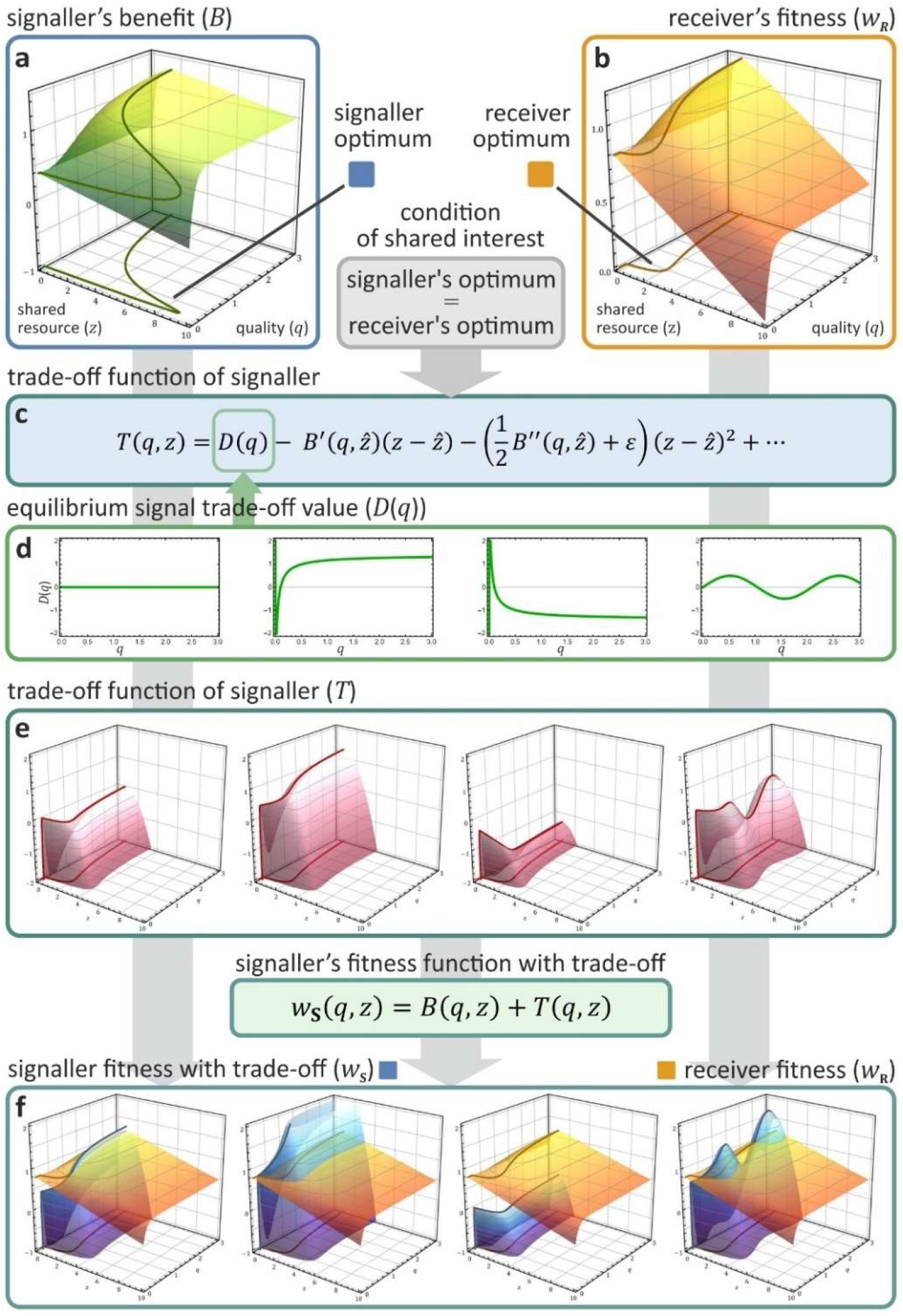
The effect of different trade-off functions on the fitness of the signaller in case of Godfray’s additive model^14^. **a**: The signaller’s benefit function *B* (without trade-off; dependent on its quality *q* and the received amount of resource *z*) defines its optimum strategy for any *q* (dark green curve; optimum curves are also projected onto the *q*-*z* baseplane for all surfaces). **b**: The receiver’s fitness function *w*_**R**_ defines its optimum strategy for any signaller quality and resource shared (yellow curve). **c**: At the honest equilibrium, the trade-off function *T* ensures that the signaller’s optimum coincides with the receiver’s optimum (for the derivation of the terms of *T*, see **Fig. 3**). **d**: An arbitrary set of equilibrium signal cost functions *D*(*q*) is selected (green curves) from left to right: 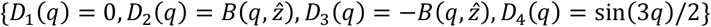, where 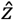 is the optimum transfer of the receiver for the given quality *q*. **e**: For any *D*_*i*_(*q*), a trade-off function *T*_*i*_ is generated (red surfaces), describing the cost value of signals in and out of equilibrium. **f**: The trade-off function *T* transforms the benefit function *B* of the signaller to the fitness function *w*_**S**_ (blue surfaces) such that its optimum strategy coincides with the receiver’s optimum strategy (yellow surfaces replicate the receiver’s fitness *w*_**R**_ as of panel **b**; note different scaling). Projected optima of *w*_**S**_ and *w*_**R**_ entirely overlap at the *q*-*z* baseplane. Parameters are {*ψ* = 1/2, *γ* = 1/2, *G* = 0.08, *U* = 1, *Z* = 10, *ε* = 1}, for details, see Appendix 2 and 4.

The second coefficient 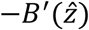 describes the *equilibrium path*, where the first derivative of *w*_**S**_ with regard to the amount of shared resource *z* is zero. This coefficient specifies that 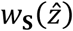 is an extremum, according to the shared-interest condition (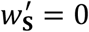, Eq.1a).

The third coefficient 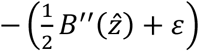 determines the steepness of the surface along the *z* dimension when deviating from the equilibrium path (*stability condition*). The condition *ε* > 0 ensures that this term is larger than the second derivative of *B* for the slope to be negative (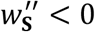, see Eq. 1b) so that 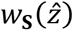 is a maximum. When this term is equal or smaller than the second derivative of *B* then the signaller’s strategy is not an equilibrium strategy. The conditions of honest signalling do not restrict higher order terms of the series therefore they can be arbitrarily chosen.

The Taylor series expansion allows functional decomposition of equilibrium trade-off (cost), equilibrium path, and stability. Accordingly, the equilibrium trade-off function *D*(*q*) can be negative, zero or even positive. These conditions can be interpreted as costly signals, cost-free signals, and signals with only benefits, respectively. For any equilibrium path, reflecting the receiver’s optimisation problem, there is an infinite number of equilibrium trade-off functions *D*(*q*) for the signaller (see **Fig. 4** for examples). In general, the *equilibrium* trade-off function (zeroth order term) is not constrained by the *equilibrium path* (first order term) or the *stability condition* (second order term).

**Fig. 4** shows four possible equilibrium signal cost functions with constant, monotonically increasing, monotonically decreasing, and oscillating trade-off functions. While the last choice seems unrealistic, it proves our point that any arbitrary, continuously differentiable function can be chosen as the equilibrium trade-off function *D*(*q*) because ***the equilibrium signal cost is independent of the stability condition*** in additive models.

### Multiplicative fitness

For multiplicative fitness, the conditions of honest signalling imply the following general form for the trade-off function *T*_*M*_ (derivatives are all evaluated at equilibrium 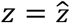 see Appendix 3 and Fig. S1 for details):

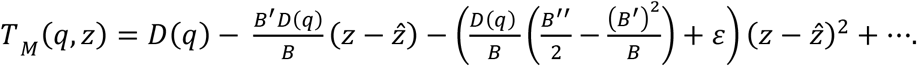

While the same functional separation is derived as in case of additive fitness functions, the same independence of terms cannot be achieved because of the multiplication of the functions. Previous models have shown that signal cost functions *D*(*q*) exist where the cost paid at the equilibrium by honest signallers is arbitrarily close to zero in multiplicative models (e.g. Lachmann *et al*.^23^ derived this result for specific cost functions in Grafen’s signalling model^12^). Our formula for *T*_*M*_ provides a method to derive all solutions for any asymmetrical signalling game with continuous (and at least twice differentiable) fitness functions. Moreover, as a novel result, it reveals, that ***equilibria with cost-free or beneficial signals exist in multiplicative models too***, not only in additive models.

In Appendix 4, we derive the general trade-off functions for well-known biological signalling games^12,15,22^. Table S1 provides a comparison of notation across these models, Table S2 compares the Taylor coefficients of the general trade-off functions of the additive and multiplicative cases, while Table S3 compares the Taylor coefficients of relevant models.

## Discussion

Our analyses provide several important results concerning the evolution of honest signals. First, we provide a general methodology for deriving the full set of an infinite number of trade-off functions for asymmetric, continuous pairwise signalling games, which allow for honest signalling. This general class of trade-off functions consists of three components: the first term defines the cost (or benefit) of a signal at the evolutionary equilibrium, the second one defines the path along the equilibrium, and the third term specifies the stability condition at the equilibrium. Second, we confirm that the results of asymmetric signalling models depend upon whether fitness effects are multiplicative or not^7^. For additive fitness, these three components are independent of each other. For multiplicative fitness models, these terms are not independent. However, the equilibrium cost of signals can be anything, zero or even negative in both fitness models and yet signalling remains honest and evolutionarily stable as long as the stability condition is fulfilled. A negative cost signal is beneficial (for an example, called the ‘Lazy Student’ game, see Appendix 8). We show the existence of such beneficial equilibria for the seminal models of the field, Grafen’s model of sexual selection^12^ and Godfray’s model of parent-offspring communication^15^. Third, these results imply that Zahavi’s Handicap Principle^3^, handicap interpretations of Grafen’s model^12^, and so-called ‘costly signalling’ models^12,15^ are incorrect in concluding that signal costs at equilibrium are a necessary condition for the evolution of honest signalling.

Our results reveal an important limitation and a surprising implication of simple asymmetric signalling games. Our model does not specify the magnitude of signal intensity at equilibrium, and just like the equilibrium signal cost, the magnitude can be any continuously differentiable function^17^. For example, in Godfray’s signalling model^15^, the equilibrium signal intensity (as a function of quality *c*) has a maximum (see Fig. 2 at^13^). Accordingly, the quality half-space below *c* = 0.5 was omitted to ignore the maximum, resulting in a monotonic decreasing signal intensity function. More generally, it was recently shown that it is possible to construct such “dishonest”, non-monotonous functions for a large class of signalling games^17^. In summary, overly simplified game-theoretical models have generated the apparent paradox that honest signalling games, which assume honesty at equilibrium, need not result in honest signalling, i.e., they need not result in a monotone increasing or decreasing signal intensity function at the equilibrium. This paradoxical result implies that the simplest possible model of honest signalling has not been sufficiently constrained in previous models, as they allow “honest” solutions where the signal cannot be used to predict the real quality of the signaller.

Existing models has another limitation, which was introduced by the use of biologically-inspired constraints^12,13,15,22^. The most common assumptions are: (*i*) the signal cost as a function of signal intensity increases monotonically (see above); and (*ii*) the equilibrium cost function *D*(*q*) is restricted by the assumption that the worst signaller gives no signal and has zero signal cost^12,13,15,22^. The first assumption directly excludes any non-monotonic cost function (like multimodal curves). The second assumption combined with the first excludes any potential cost function with zero or negative equilibrium signal cost. We call these the *standard costly signalling assumptions*. When these assumptions are employed, the result is the traditional “costly signalling” outcome in which the equilibrium cost function has positive values with monotonically increasing signal cost (i.e. equilibrium signals are honest and costly, see e.g., Grafen’s strategic choice signalling model^12^). In other words, the central handicap claim that signals must be costly (or wasteful) at the honest equilibrium does not follow from the most general formulation of the conditions of honest signalling; it follows only from the additional constraints – the *standard costly signalling assumptions*. Note that the problem here is not the application of specific assumptions, but rather the misinterpretation that signal cost directly follows from the general formulation (e.g., Grafen asserted that “If we see a character which does signal quality, then it must be a handicap”^12^ p. 521). These costly signalling assumptions may or may not be realistic, but the interpretation that costly equilibrium is a necessity is incorrect, as it does not follow from the general formulation. This misinterpretation of Grafen’s model is what led to the widespread acceptance of the Handicap Principle and the popularity of ‘costly signalling theory’ (for a detailed discussion of misinterpretations, see^5^).

It is important to note that our findings also highlight the limitations of studying the evolutionary equilibrium for honesty using game theoretical models. Our formula clears up confusion over differences between conclusions that follow from the *conditions* of honest signalling models versus the consequences from *additional constraints*. We show that honest signalling models can only predict the value of marginal change – the behaviour of the system – in vicinity of the evolutionary equilibrium (by definition) without including additional biological constraints. They cannot provide predictions about (*i*) the cost of signals at or outside of the equilibrium, or (*ii*) the marginal change further away from the equilibrium path (see **Fig. 4**). One can add additional constraints to the models (see above discussion, e.g.^12,15^) but then the results are simply determined by these constraints. When postulates (standard costly assumptions) unnecessarily constrain a model (honest signalling), the conclusion (honest signals are costly) may be true not because of the model but because of the postulates. Making such general conclusions and ignoring a model’s limitations due to its restrictive assumptions is an example of the *overgeneralization fallacy*. We have demonstrated here how removing the constraints of these previous models undermines their usual interpretations: honest signals need not be costly (see **Fig. 5**), and hence the Handicap Principle can be fully rejected. If honest signals turn out to be costly in nature, such results do not explain their evolution, regardless of the magnitude of their costs.

**Fig. 5.**
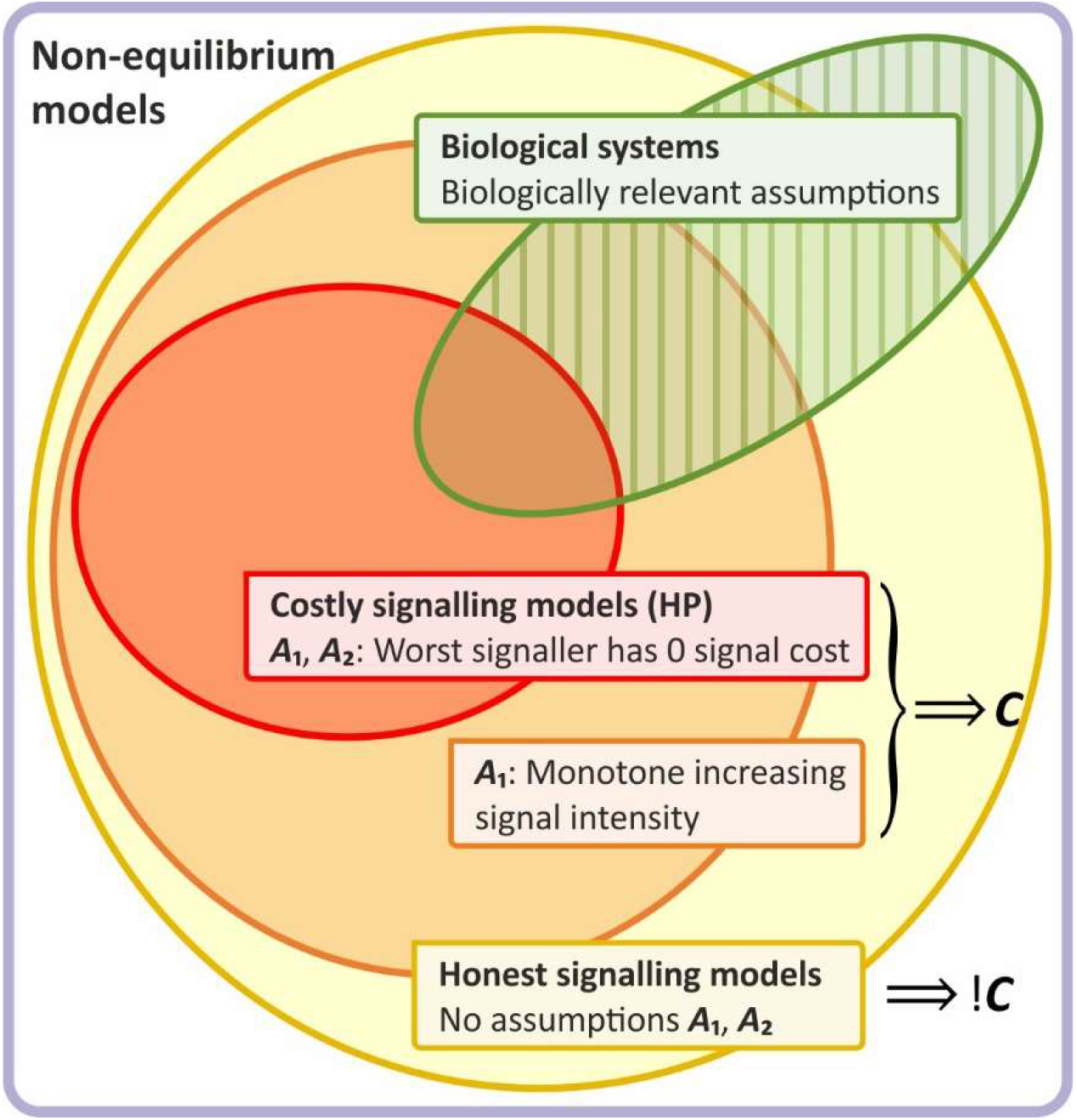
The overgeneralization fallacy of the Handicap Principle (HP). When postulates unnecessarily constrain a model, the conclusion may be true not because of the model (i.e., because of more general properties), but because of the postulates. When one nevertheless claims that the conclusion is generally true regardless of postulated assumptions, that is a fallacy of overgeneralization. If removing the constraints switches the conclusion (strongly depending thus on the assumptions), one has also committed a category mistake, incorrectly attributing properties to the model and missing its true nature. Standard costly signalling assumptions (SCSA = ***A***_1_ & ***A***_2_, red set) unnecessarily constrain the model of honest signalling (yellow set), because they exclude a potentially important class of cost functions. Moreover, biologically relevant assumptions may not be constrained to SCSA, contrary to what Grafen suggests (all biological signals are handicaps, implying that the green set is a subset of the red set, and the striped subset is empty)^12^; see Discussion). Removing SCSA switches the conclusion ***C*** of the model to ! ***C***: honest signals need not be costly, as we have proved in this paper. Hence the generalization that the conclusion of the model is valid without SCSA is wrong. That is, the conclusion of the Handicap Principle (***C*** = honest signals are costly), stems from its assumptions and not from more general properties, leading to tautology and a category mistake. Even if honest signals turn out to be costly in nature, the Handicap Principle cannot account for them.

Results of our model help to explain why empirical studies do not match the predictions generated by the Handicap Principle or costly signalling theory. Numerous studies have failed to find “costly signals” predicted by Zahavi and Grafen (see^45–52^; for example, offspring begging calls are nowhere as costly as often assumed^45^, see^46^ for review). Not even the elaborate peacock’s train, the flagship example of the Handicap Principle, fits the predictions of costly signalling models. The train does not hinder movement^47^; on the contrary, males with longer trains are able to take-off faster than males with shorter ones^48^. Empirical studies have often been cautious with their conclusions and suggested that some other types of signalling costs might be discovered that would support the Handicap Principle. Our results show that such signal costs are neither sufficient nor necessary to explain honesty; **they are simply irrelevant for signal honesty**. Cost-free or even beneficial honest equilibria are possible: high-quality offspring need not waste energy to produce honest begging calls and peacock trains need not be wasteful or even costly to signal male quality. **Honesty is maintained by the *potential costs* of cheating through dishonest signals, but not by some general cost of signalling at the evolutionary equilibrium** (see^6,23^). It follows that many efforts to measure signalling cost at the equilibrium are not informative about honesty or stability of the signalling system at all. While similar arguments previously have been made^6,23^, our equations provide the first mathematical proof.

Traditional explanations of honesty differentiate between cost (i.e. ‘handicaps’) and constraints (i.e. ‘indices’). Recent modelling work seems to undermine such strict dichotomy. First, these models suggest a continuum from cues to costly signals (see^53^), second, they claim “that costly signalling theory provides the ultimate, adaptive rationale for honest signalling, whereas the index hypothesis describes one proximate (and potentially very general) mechanism for achieving honesty.” (see^54^, abstract). While we agree that physical constraints pose a potential proximate mechanism our results do not support the first part of the claim, namely that “costly signalling theory provides the ultimate, adaptive explanation for honest signalling…” (^54^). While we also agree with the claim that there is a continuum from cheap to more costly signals, there is also potential continuum from cost-free to beneficial signals, which was ignored by costly signalling theory.

In summary, costly signalling models have unnecessarily restricted the set of possible solutions to explain the evolution of honesty. Consequently, the costly signalling solution in traditional costly signalling models is not so unexpected or interesting, being the consequence of the very assumptions postulated by these models. Due to their restricted design, these models could only investigate costly signalling solutions because any other solution (e.g., cost-free or beneficial) is impossible within their boundaries due to the additional *costly signalling assumptions*. Given their restrictive set of assumptions, these models consequently cannot provide general results for honest signalling or general predictions about the evolution of signals (**Fig. 5**). Our method provides a general calculation of signalling equilibria therefore, it should allow the field of honest signalling theory to progress beyond the domain of restricted costly signalling models.

These results highlight the need for a better framework than the Handicap Principle (or costly signalling) for explaining honest signalling. Signals are expected to confer fitness trade-offs (*signalling trade-offs)*, which are better understood as life-history trade-offs rather than as ‘handicaps’ (i.e. signals that are honest because they are costly). All of the seminal models of the Handicap Principle used life-history trade-offs to create differential cost between signallers of different quality: a trade-off between reproduction and survival^12^ or a trade-off between current and future offspring^15^. A recent laboratory experiment shows the importance of condition-dependent trade-offs versus equilibrium costs of signalling^55^. A recent model investigated the differences between additive and multiplicative fitness functions also adopts an evolutionary trade-off framework^56^. Our results here show how honesty can be selectively maintained by condition-dependent signalling trade-offs. Such trade-offs can be difficult to measure^57,58^, but this approach allows the use of theoretical models and empirical methodology established in this field^58–61^.

Our results highlight the critical failure of the Handicap Principle to understand the mechanism of honest signalling. Cost-free or even beneficial honest equilibria can exist because honest signals need not be wasteful. On the contrary, honest signallers can be (and will be) ***honest and efficient***, it is the potential cheater that must be ***less efficient*** (more wasteful) than honest signallers. This optimality or efficiency principle is not new (see^25^), but unfortunately it has been overshadowed by the erroneous Handicap Principle. The efficiency principle can explain the existence of the seemingly exaggerated features like the train of the peacock or the antlers of deer. Trains and antlers can be extravagant, yet provide minimal or no fitness cost to the bearer^52^ because honest signallers can produce them efficiently. The evolution of honest signals does not fall under a separate principle as asserted by Zahavi, as signals evolve according to the same evolutionary processes as any other trait: *through natural selection, which favours efficiency over wastefulness*.

## Methods

### The model

The model consists of two agents, the signaller **S** and the receiver **R**. The signaller elicits a signal to request an amount *a* of the resource from the receiver. The signaller’s fitness *w*_**S**_ depends on the signaller’s quality *q*, on the intensity of its signal (asking for an amount *a* of the resource) and on the amount of resource *z* provided by the receiver due to the signal. The receiver’s fitness *w*_**R**_ depends on the hidden quality *q* of the signaller and on its own response strategy, that specifies the amount of resource *z* the receiver shares with the signaller; 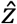 denotes the equilibrium amount (in honest equilibrium, it is expected that 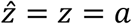). The response can be written as a function directly dependent on *q*. We treat the signaller fitness *w*_**S**_ as an additive or multiplicative combination of a benefit function *B* and a signal trade-off function *T*, where both *B*(*q, z*) and *T*(*q, z*) are functions of signaller quality *q* and the received resource *z. T* defines the trade-off of asking for *z* = *a* amount of resource as a signaller of quality *q*, depending entirely on the signaller. *B*, on the other hand, is controlled entirely by the receiver’s response (how much *z* the receiver shares, based indirectly, through a signal, on the signaller’s quality *q*; see Appendix 1 for a formal derivation). This interpretation justifies the mathematical decomposition of *w*_**S**_ into these two functions. Derivatives are with respect to *z*; a hat over a symbol indicates equilibrium value. **Table 1** lists the quantities of the model.

**Table 1.**
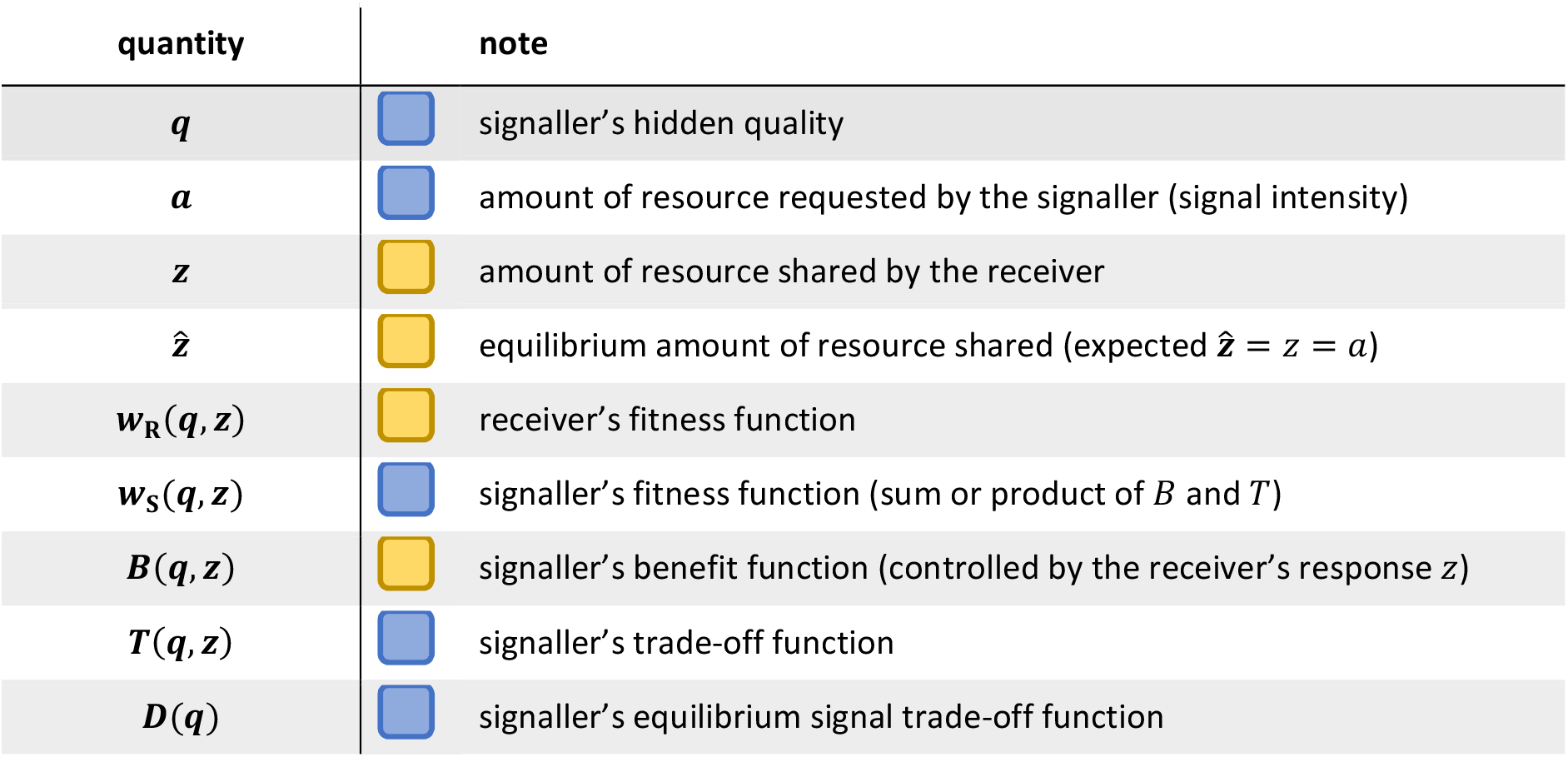
Notation used in the model. Coloured boxes indicate the controlling party, blue for signaller, yellow for receiver. For more details, see Appendix 1 Table S2.

| quantity | note |  |
| --- | --- | --- |
| $q$ | | signaller's hidden quality |
| $a$ | | amount of resource requested by the signaller (signal intensity) |
| $z$ | | amount of resource shared by the receiver |
| $\hat{z}$ | | equilibrium amount of resource shared (expected $\hat{z} = z = a$) |
| $w_R(q, z)$ | | receiver's fitness function |
| $w_S(q, z)$ | | signaller's fitness function (sum or product of $B$ and $T$) |
| $B(q, z)$ | | signaller's benefit function (controlled by the receiver's response $z$) |
| $T(q, z)$ | | signaller's trade-off function |
| $D(q)$ | | signaller's equilibrium signal trade-off function |

### Conditions of honest equilibrium

The honest signalling equilibrium has two conditions (for details, see Appendix 1):

1. *Condition of honest optimum* specifies that there exists an optimum amount of resource 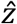 that the receiver is willing to share. That is, the receiver, depending on the received signal, shares an amount that equals to the amount it would share if he could directly assess the signaller’s quality. This means, that signals are honest as they reveal the signaller’s quality, so that resource allocation is optimal for the receiver.
2. *Condition of shared interest* specifies that there is no conflict between receiver and signaller as the signaller asks the exact amount the receiver is willing to share and both *w*_**S**_ *and w*_**R**_ have their respective maxima at 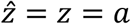. Since neither receiver nor signaller wants to deviate from it, the condition implies stability. It has two conditions:
  2.a *Extremum condition* specifies that *w*_**S**_ has an extremum at 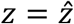:

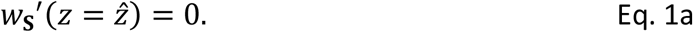
  2.b *Stability condition* specifies that the extremum at 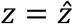 is a maximum:

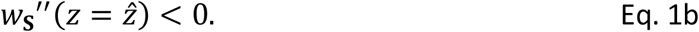

### Reverse-engineering the trade-off function

Finding honest signalling solutions in signalling games requires: (*i*) calculating the optimal resource sharing decision for the receiver, (*ii*) calculating signalling trade-offs (*T*, traditionally called ‘cost function’) that transform the signaller’s optimal decision to the receiver’s optimum. That is, the signaller has optimal fitness when it asks for and gets the same amount *z* the receiver is willing to share in its fitness optimum (see **Fig. 1**). We provide a formal method to reverse engineer the trade-off function *T* that is general and specifies all the infinite number of solutions. We use a Taylor series expansion of the signaller’s fitness *w*_**S**_ to specify the conditions of honest signalling identified by previous models^13,22^. Since *w*_**S**_ is composed of *B* and *T* (additively or multiplicatively), *w*_**S**_ can be expressed as the appropriate combination of terms of the Taylor series of *B* and *T* (see **Fig. 3** and Fig. S1). The first and second order Taylor coefficients of *w*_**S**_ can be used to express the stability and honesty conditions (Eq. 1a, b) as constraints on the first and second derivatives of *B* and *T*. Since *B* is given, we use these constraints to construct a general trade-off function *T* that, when combined with *B*, yields a signaller fitness function *w*_**S**_ that fulfils the conditions of honest signalling (its optimum coincides with the optimum of *w*_**R**_). In Appendix 4, we apply our method to known models.

### Additive fitness model

First, we derive the general trade-off function for the additive model. For a visual guide, see the left panel of Fig. S1, for details, see Appendix 2. In case of the additive fitness model, the signaller’s fitness is the sum of the benefit and trade-off functions:

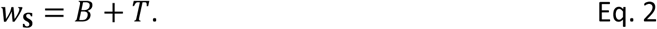

Both *B* and *T* can be written as Taylor series around the equilibrium 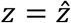 (omitting function arguments *q* and *z* for sake of simplicity):

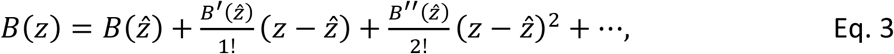

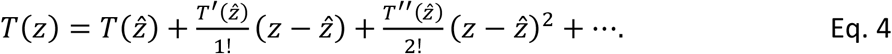

With *β*_*k*_ = *B*^(*k*)^/*k*! and *τ*_*k*_ = *T*^(*k*)^/*k*!, the sum of *B* and *T* can be rewritten:

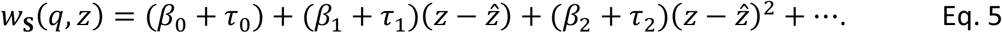

At the equilibrium 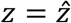, conditions Eqs. 1a, b must be met by *w*_**S**_. According to Eq. 1a, the first derivative of *w*_**S**_ must be zero:

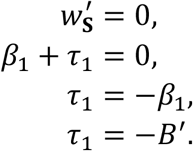

According to Eq. 1b, the second derivative of *w*_**S**_ must be smaller than zero:

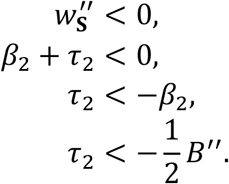

The inequality is always satisfied if *ε* > 0:

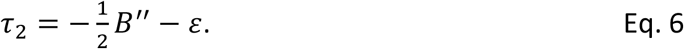

Substituting *τ*_1_ and *τ*_2_ into Eq. 4 and *D*(*q*) for 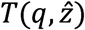, the constructed trade-off function for additive fitness components is:

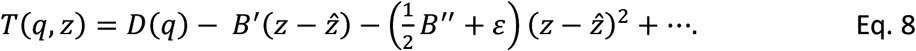

### Multiplicative fitness model

In case of the multiplicative fitness model, the signaller’s fitness is the product of the benefit and trade-off functions (for a visual guide, see the right panel of Fig. S1, for details, see Appendix 3):

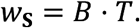

The Taylor series of a multiplicative *w*_**S**_ is the product of the individual Taylor series of the composite functions *B* and *T* (Eqs. 3, 4):

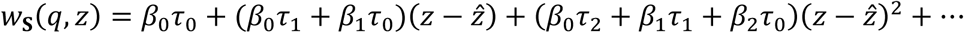

At the equilibrium 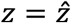, conditions Eqs. 1a, b must be met by *w*_**S**_. According to Eq. 1a, the first derivative of *w*_**S**_ must be zero (omitting function arguments):

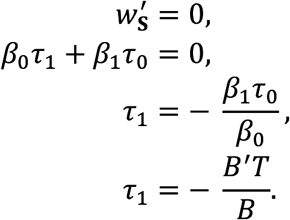

The first derivative of *T* at the equilibrium depends on *T* itself, unlike in the additive case. According to Eq. 1b, the second derivative of *w*_**S**_ must be smaller than zero (substituting *τ*_1_ from above):

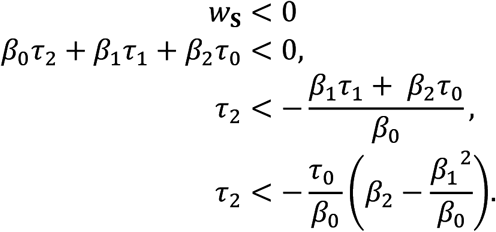

The inequality is always satisfied if *ε* > 0:

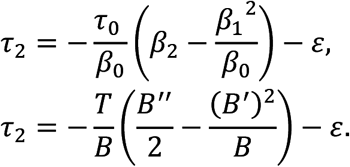

Substituting *τ*_1_ and *τ*_2_ into Eq. 4 and *D*(*q*) for 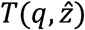, the constructed trade-off function for multiplicative fitness components is:

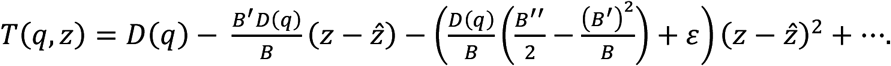

## Acknowledgements & funding

The authors acknowledge support from the Hungarian Research Fund under grant numbers NKFIH #132250 (SS), #140901 (IZ), #129848 (DC, IZ), from the Templeton World Charity Foundation TWCF0268 (DC), from the Austrian Science Fund P28141-B25 (DJP) and from the Human Frontier Science Program RGP003/2020 (DJP).

## Author contributions

SS, DC and DJP conceived the idea, SS, DC and IZ designed and analysed the model, IZ and SS designed and created the figures, SS, IZ, DC and DJP wrote the paper. All authors contributed to the article and approved the submitted version.

