## supplementary materials for "General solution to biological signalling games: costly signalling and beyond"

10 May 2022

Szabolcs Számadó[1,2,§*], István Zachar[3,4,5,§], Dániel Czégel[3,5,6,7] and Dustin J. Penn[8]

[1] Department of Sociology and Communication, Budapest University of Technology and Economics, Egry J. u. 1. H-1111 Budapest, Hungary

[2] CSS-RECENS „Lendület" Research Group, MTA Centre for Social Science, Tóth Kálmán u. 4., H-1097 Budapest, Hungary

[3] Institute of Evolution, MTA Centre for Ecological Research, Konkoly-Thege Miklós út 29-33., H-1121 Budapest, Hungary

[4] MTA-ELTE Theoretical Biology and Evolutionary Ecology Research Group, Eötvös Loránd University, Department of Plant Taxonomy and Ecology, Pázmány Péter sétány 1/c, H-1117 Budapest, Hungary

[5] Parmenides Foundation, Centre for the Conceptual Foundation of Science, Kirchplatz 1., D-82049 Pullach im Isartal, Germany

[6] Doctoral School of Biology, Institute of Biology, Eötvös Loránd University, Pázmány Péter sétány 1/C, H-1117 Budapest, Hungary

[7] BEYOND Center for Fundamental Concepts in Science, Arizona State University, AZ 85287–0506 Tempe, Arizona, USA

[8] Department of Interdisciplinary Life Sciences, Konrad Lorenz Institute of Ethology, University of Veterinary Medicine, Vienna, Savoynestrasse 1a, 1160 Vienna, Austria

[§] Shared first authorship

#### Appendix 1. Conditions of honest signalling

The model consists of two agents, the signaller $\mathbf{S}$ and the receiver $\mathbf{R}$. The signaller elicits a signal of intensity $x$ to request an amount $a$ of the resource from the receiver. The signaller's fitness $w_{\mathbf{S}}$ depends on the signaller's quality $q$, on the intensity of its signal $x$ (asking for an amount $a$ of the resource) and on the amount of resource $z$ provided by the receiver due to the signal $x$:

$$x = S_{\mathbf{S}}(q),$$
$$w_{\mathbf{S}}(q, x, z) = w_{\mathbf{S}}(q, S_{\mathbf{S}}(q), z).$$

which can be simplified to depend on $q$ and $z$, as $w_{\mathbf{S}}(q, z)$.

The receiver's fitness $w_{\mathbf{R}}$ depends on the signal intensity $x$ of the signaller (thus indirectly, on its hidden quality $q$) and on its own response strategy $S_{\mathbf{R}}$, that specifies the amount of resource $z$ the receiver shares with the signaller.

$$z_x = S_{\mathbf{R}}(x) = S_{\mathbf{R}}\big(S_{\mathbf{S}}(q)\big),$$
$$w_{\mathbf{R}}(x, z) = w_{\mathbf{R}}\big(x, S_{\mathbf{R}}(x)\big) = w_{\mathbf{R}}\Big(x, S_{\mathbf{R}}\big(S_{\mathbf{S}}(q)\big)\Big).$$

Alternatively, the response can be written as the strategy $Z_{\mathbf{R}}$ directly dependent on $q$:

$$z_q = Z_{\mathbf{R}}(q)$$

We treat the signaller fitness $w_{\mathbf{S}}(q, z)$ as an additive or multiplicative combination of a benefit function $B(q, z)$ and a signal trade-off function $T(q, z)$:

| additive fitness | multiplicative fitness |
|---|---|
| $w_{\mathbf{S}}(q, z) = B(q, z) + T(q, z)$ | $w_{\mathbf{S}}(q, z) = B(q, z) \cdot T(q, z)$ |

$B$ is controlled by the receiver's response strategy $S_{\mathbf{R}}(x) = z$:

$$B(q, z) = B\big(q, S_{\mathbf{R}}(x)\big).$$

$T$, on the other hand, depends entirely on the signaller, defining the cost of asking for $z = a$ amount of resource as a signaller of quality $q$:

$$T(q, \mathrm{z} = a) = T\big(q, S_{\mathbf{S}}(q)\big).$$

This interpretation justifies the mathematical decomposition of $w_{\mathbf{S}}$ into these two functions. Throughout the mathematical analysis, derivatives are with respect to $z$; a hat over a symbol indicates equilibrium value. Table S1 lists the quantities of the model.

*Table S1. Comparison of notation of certain relevant models. G90:* (Grafen 1990)*, G91:* (Godfray 1991)*, NS99:* (Nöldeke and Samuelson 1999)*, BSL02:* (Bergstrom et al. 2002)*; SCZ19:* (Számadó et al. 2019).

| | **G90** | **G91** | **NS99** | **BSL02** | **SCZ19** | **This model** |
|---|---|---|---|---|---|---|
| signaller | | | | | | $\mathbf{S}$ |
| signal receiver (listener) | | | | | | $\mathbf{R}$ |
| hidden quality of the signaller (condition, trait) | $q$ | $c$ | $c$ | $q$ | $c$ | $q$ |
| amount of resource the signaller is asking for | | $x$ | | | | $a$ |
| strategy of the signaller (signal intensity) | $a = A(q)$ | $x$ | $x$ | $S$ | $x$ | $x = S_{\mathbf{S}}(q)$ |
| amount of resource shared by the receiver (signaller gets this much) | | $y$ | | | | $z$ |
| strategy of the receiver depending on signal intensity | $p = P(a)$ | $y = S(x)$ | $z$ | $R$ | $z$ | $z = S_{\mathbf{R}}(x)$ |
| strategy of the receiver depending directly on signaller's quality | | $y$ | $z$ | $R$ | $z$ | $z = Z_{\mathbf{R}}(q)$ |
| fitness of receiver | | $f(y,x,c)$ $+ g(y)$ | $u$ | $G$ | $w_{\mathbf{R}}$ | $w_{\mathbf{R}}$ |
| fitness of signaller | $w(a,p,q)$ | $f(y,x,c)$ | $v$ | $H$ | $w_S$ | $w_{\mathbf{S}} = B + T$ or $w_{\mathbf{S}} = B \cdot T$ |
| benefit of signaller | $w(a,p,q)$ | $f(y,x,c)$ | $v$ | $H$ | $w_S$ | $B$ |
| signal cost of signaller as a function of signal intensity | | | $f(x)$ | | $f(x)$ | $f(x)$ or $f(q,x)$ |
| signal cost of signaller as a function of quality and the strategy of the receiver | | | $L(c,z)$ | $c(q,R)$ | $L(c,z)$ | $T(q,z) = f\left(q, S_{\mathbf{R}}^{-1}(z)\right)$ or more appropriately: $T(q,x) = T\left(q, S_{\mathbf{S}}(a)\right)$ |
| equilibrium signal cost as a function of signaller quality | | | | | | $D(q)$ |
| **IDENTITIES** | | | | | amount of resource shared in equilibrium | $S_{\mathbf{S}}(a) = S_{\mathbf{R}}^{-1}(x) = \hat{z}$ |
| | | | | | in equilibrium | $S_{\mathbf{R}}^{-1}(z) = S_{\mathbf{R}}(x)$ |

Honest equilibrium has two conditions:

1. *Condition of honest optimum*: When this condition is met, signals are honest as they reveal the signaller's quality, so that the receiver can respond adaptively. Accordingly, the equilibrium strategy of the receiver $S_{\mathbf{R}}(x)$ depending on the signaller's signal $x$ is the same as the strategy $Z_{\mathbf{R}}(q)$ if receiver would be able to directly assess the signaller's quality:

$$\hat{z} = \hat{S}_{\mathbf{R}}(\hat{x}) = \hat{Z}_{\mathbf{R}}(q). \qquad \text{Eq. S1}$$

That is, the equilibrium signal $\hat{x}$ honestly conveys the quality $q$ of the signaller, so the receiver shares an optimum amount of resource $\hat{z}$ with a signaller of quality $q$ using signal $\hat{x}$. Consequently, the signaller receives a resource allocation that is optimal for the receiver.

2. *Condition of shared interest*: When this condition is met, there is no conflict between receiver and signaller as the signaller asks the exact amount the receiver is willing to share. That is, at an honest equilibrium, both the signaller's and receiver's fitness optimization problems for their respective fitness functions ($w_{\mathbf{R}}$ and $w_{\mathbf{S}}$) yield the same resource allocation, $\hat{z}$:

$$\hat{a} = \arg\max\big(w_{\mathbf{R}}(z)\big),$$
$$\hat{z} = \arg\max\big(w_{\mathbf{S}}(z)\big),$$
$$\hat{a} = \hat{z}, \qquad \text{Eq. S2}$$

that is, both $w_{\mathbf{S}}$ *and* $w_{\mathbf{R}}$ have their respective maxima at $\hat{z}$. This condition implies stability, since neither the receiver nor the signaller wants to deviate from it, and itself has two conditions:

*2.a. Extremum condition:* the first derivative of the signaller's fitness as the function of the allocated resource $z$ is zero at $\hat{z} = \hat{Z}_{\mathbf{R}}(q)$ (called the *equilibrium path* at (Bergstrom et al. 2002)):

$$w_{\mathbf{S}}'\left(\hat{Z}_{\mathbf{R}}(q)\right) = 0,$$

or simply:

$$w_{\mathbf{S}}'(z = \hat{z}) = 0. \qquad \text{Eq. S3}$$

*2.b. Stability condition: t*he second derivative of the signaller's fitness as the function of the allocated resource $z$ is smaller than zero at $\hat{z} = \hat{Z}_{\mathbf{R}}(q)$:

$$w_{\mathbf{S}}''\left(\hat{Z}_{\mathbf{R}}(q)\right) < 0,$$

or simply:

$$w_{\mathbf{S}}''(z = \hat{z}) < 0. \qquad \text{Eq. S4}$$

### Appendix 2. Derivation of the general trade-off function for the additive model

First, we derive the general trade-off function for the additive model. For a visual guide, see the left panel of Fig. S1. In case of the additive fitness model, the signaller's fitness is the sum of the benefit and trade-off functions:

$$w_{\mathbf{S}} = B + T. \qquad \text{Eq. S5}$$

The signaller benefit function $B$ can be written as a Taylor series around $z = \hat{z}$.

$$B(q,z) = B(q,\hat{z}) + \frac{B'(q,\hat{z})}{1!}(z-\hat{z}) + \frac{B''(q,\hat{z})}{2!}(z-\hat{z})^2 + \cdots = \sum_{k=0}^{\infty}\frac{B^{(k)}(q,\hat{z})}{k!}(z-\hat{z})^k,$$

where $B^{(k)}$ denotes the $k^{th}$ derivative of $B$ with respect to $z$. We can omit function arguments for sake of simplicity:

$$B(q,z) = \sum_{k=0}^{\infty} \frac{B^{(k)}}{k!}(z-\hat{z})^k = B + B'(z-\hat{z}) + \frac{B''}{2!}(z-\hat{z})^2 + \cdots. \qquad \text{Eq. S6}$$

Similarly, the Taylor expansion of the signal trade-off function $T$ around $\hat{z}$ is:

$$T(q,z) = \sum_{k=0}^{\infty} \frac{T^{(k)}}{k!}(z-\hat{z})^k = T + T'(z-\hat{z}) + \frac{T''}{2!}(z-\hat{z})^2 + \cdots. \qquad \text{Eq. S7}$$

Let's introduce shorthand notations $\beta_i$ and $\tau_i$ for the $i^{\text{th}}$ Taylor coefficients of $B$ and $T$, respectively:

$$\beta_k = \frac{B^{(k)}}{k!} \quad \longrightarrow \quad B(q,z) = \beta_0 + \beta_1(z-\hat{z}) + \beta_2(z-\hat{z})^2 + \beta_3(z-\hat{z})^3 + \cdots, \qquad \text{Eq. S8}$$

$$\tau_k = \frac{T^{(k)}}{k!} \quad \longrightarrow \quad T(q,z) = \tau_0 + \tau_1(z-\hat{z}) + \tau_2(z-\hat{z})^2 + \tau_3(z-\hat{z})^3 + \cdots. \qquad \text{Eq. S9}$$

In equilibrium $z = \hat{z}$, the amount of resource requested equals the amount shared by the receiver (Eq. S2), hence the cost of signalling depends entirely on the quality of the signaller. Therefore, we identify $\tau_0 = T(q,\hat{z})$ with the *equilibrium signal cost function* $D(q)$ that represents the cost that signallers pay at the equilibrium $z = \hat{z}$ for giving signals. At the equilibrium, $T(q,\hat{z})$ does not depend on $z$, therefore the second argument can be ignored.

$$\tau_0 = T(q,\hat{z}) = D(q), \qquad \text{Eq. S10}$$

where $D(q)$ can be any function of offspring quality $q$.

In additive models, the signaller's fitness $w_S$ is the sum of the benefit function $B$ and the trade-off function $T$:

$$\begin{aligned} w_S(q,z) &= \beta_0 + \beta_1(z-\hat{z}) + \beta_2(z-\hat{z})^2 + \tau_0 + \tau_1(z-\hat{z}) + \tau_2(z-\hat{z})^2 + \cdots = \\ &= (\beta_0 + \tau_0) + (\beta_1 + \tau_1)(z-\hat{z}) + (\beta_2 + \tau_2)(z-\hat{z})^2 + \cdots \end{aligned}$$

In equilibrium $z = \hat{z}$, the conditions Eq. S3 and Eq. S4 must be met. According to Eq. S3, the first derivative of $w_S$ must be zero at $z = \hat{z}$:

$$\begin{aligned} \frac{\partial}{\partial z} w_S(q,z) &= 0, \\ \beta_1 + \tau_1 &= 0, \\ \tau_1 &= -\beta_1. \end{aligned}$$

Substituting in $\beta_1$ from Eq. S8 gives the value of $\tau_1$ that satisfies the condition:

$$\tau_1 = -B'(q,z). \qquad \text{Eq. S11}$$

According to Eq. S4, the second derivative of $w_S$ must be smaller than zero at the equilibrium:

$$\begin{aligned} \frac{\partial^2}{\partial z^2} w_S(q,z) &< 0, \\ (\beta_2 + \tau_2) &< 0, \\ \tau_2 &< -\beta_2, \\ \tau_2 &< -\frac{1}{2} B''(q,z). \end{aligned}$$

The inequality is always satisfied if $\varepsilon > 0$:

$$\tau_2 = -\frac{1}{2} B''(q,z) - \varepsilon. \qquad \text{Eq. S12}$$

The general form of any, twice differentiable, equilibrium trade-off function for additive fitness functions can be written by substituting $D(q)$, $\tau_1$ and $\tau_2$ from Eq. S10, Eq. S11 and Eq. S12 into Eq. S9:

$$T_A = T(q,z) = D(q) - B'(q,z)(z-\hat{z}) - \left(\frac{1}{2}B''(q,z) + \varepsilon\right)(z-\hat{z})^2 + \cdots. \quad \text{Eq. S13}$$

Higher order terms $\tau_3, \tau_4$, ... can be anything and won't affect the solution.

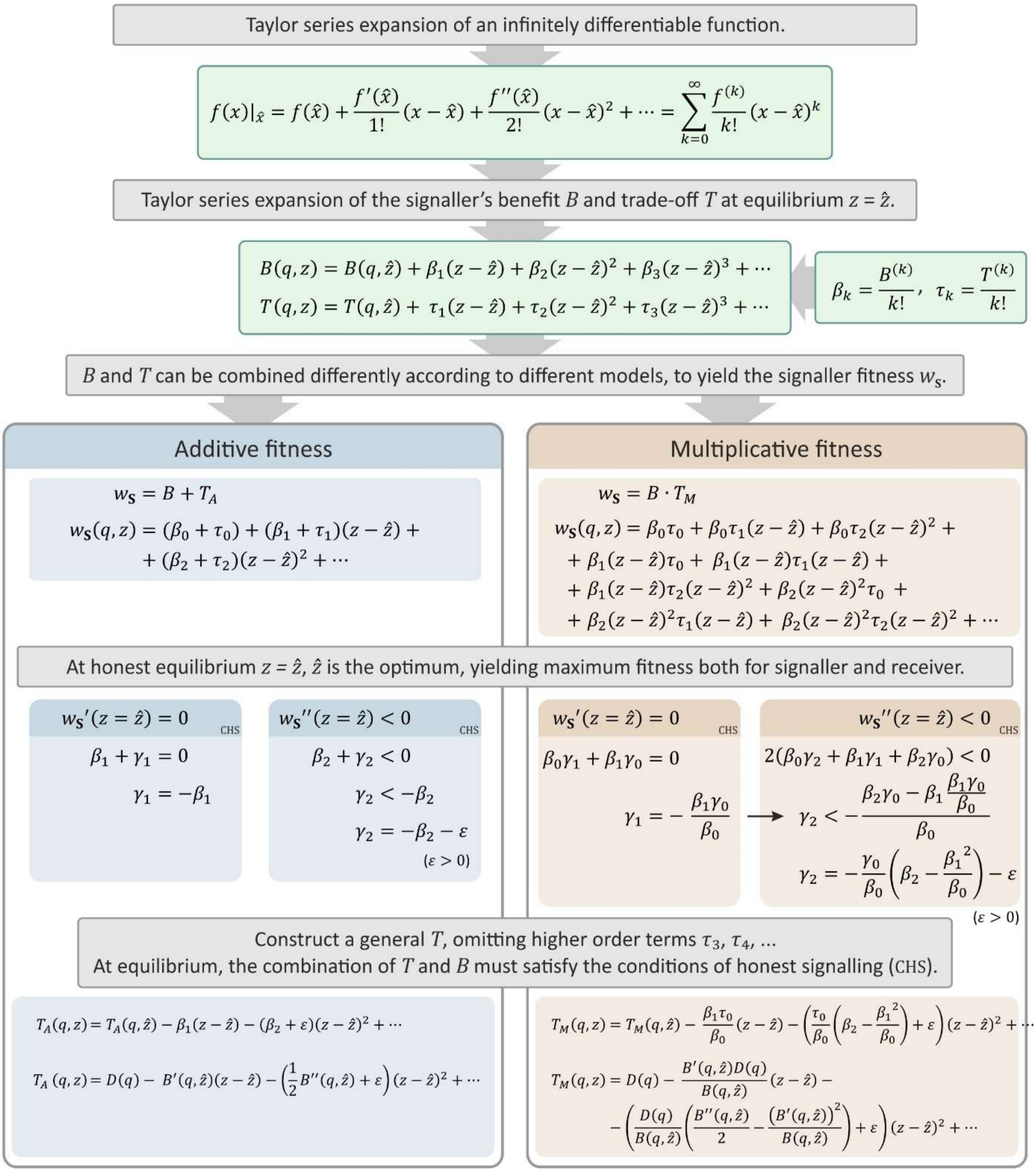


*Fig. S1. Description of the reverse-engineering method to find a cost function $D$ and to construct a trade-off function $T$ that can be combined with the benefit function $B$ into the signaller fitness $w_S$ that satisfies the conditions of honest signalling (CHS), **regardless of the equilibrium cost paid by an honest signaller**. Derivatives, when indicated with primes, are with respect to z.*

#### Appendix 3. Derivation of the general trade-off function for the multiplicative model

We follow the above outlined method to derive the general trade-off function for the multiplicative model (for a visual guide, see the right panel of Fig. S1). In case of the multiplicative fitness model, the signaller's fitness is the product of the benefit and trade-off functions:

$$w_{\mathrm{S}} = B \cdot T. \qquad \text{Eq. S14}$$

The Taylor series of a multiplicative function is the product of the individual Taylor series of the composite functions. Thus, when $B$ and $T$ are multiplied in $w_{\mathrm{S}}$, the Taylor series expansion of the product is as follows (using the shorthand notation of Eq. S8 and Eq. S9 and omitting higher order terms):

$$w_{\mathrm{S}}(q,z) = (\beta_0 + \beta_1(z-\hat{z}) + \beta_2(z-\hat{z})^2)(\tau_0 + \tau_1(z-\hat{z}) + \tau_2(z-\hat{z})^2) + \cdots.$$

After rearranging:

$$\begin{aligned} w_{\mathrm{S}}(q,z) = {} & \beta_0\tau_0 + \beta_0\tau_1(z-\hat{z}) + \beta_0\tau_2(z-\hat{z})^2 + \beta_1(z-\hat{z})\tau_0 + \beta_1(z-\hat{z})\tau_1(z-\hat{z}) + \\ & +\beta_1(z-\hat{z})\tau_2(z-\hat{z})^2 + \beta_2(z-\hat{z})^2\tau_0 + \\ & +\beta_2(z-\hat{z})^2\tau_1(z-\hat{z}) + \beta_2(z-\hat{z})^2\tau_2(z-\hat{z})^2 + \cdots. \end{aligned}$$

In equilibrium, where $z = \hat{z}$, the conditions Eq. S3 and Eq. S4 must be met. According to the stability condition Eq. S3, in the equilibrium $z = \hat{z}$, the first derivative of $w_{\mathrm{S}}$ must be zero:

$$\frac{\partial}{\partial z} w_{\mathrm{S}}(q,z) = 0,$$

$$\beta_0\tau_1 + \beta_1\tau_0 = 0,$$

$$\tau_1 = -\frac{\beta_1\tau_0}{\beta_0}. \qquad \text{Eq. S15}$$

Substituting in $\beta_0$, $\beta_1$ and $\tau_0$ gives the value of $\tau_1$ that satisfies the condition:

$$\tau_1 = -\frac{B' T}{B}. \qquad \text{Eq. S16}$$

Thus, unlike in the additive case, the first derivative of the trade-off function $T$ at the equilibrium depends on $T$ itself. According to Eq. S4, the second derivative of $w_{\mathrm{S}}$ must be smaller than zero in the equilibrium:

$$\frac{\partial^2}{\partial z^2} w_{\mathrm{S}}(q,z) < 0,$$

$$(\beta_0\tau_2 + \beta_1\tau_1 + \beta_2\tau_0) < 0,$$

$$\tau_2 < -\frac{\beta_1\tau_1 + \beta_2\tau_0}{\beta_0}.$$

After substituting $\tau_1$ from Eq. S15 and rearrangement:

$$\tau_2 < -\frac{\beta_2\tau_0 - \beta_1\frac{\beta_1\tau_0}{\beta_0}}{\beta_0},$$

$$\tau_2 < -\frac{\tau_0}{\beta_0}\left(\beta_2 - \frac{{\beta_1}^2}{\beta_0}\right).$$

The inequality is always satisfied if $\varepsilon > 0$:

$$\tau_2 = -\frac{\tau_0}{\beta_0}\left(\beta_2 - \frac{{\beta_1}^2}{\beta_0}\right) - \varepsilon. \qquad \text{Eq. S17}$$

Substituting appropriate values for $\beta$ and $\tau$ from Eq. S8 and Eq. S9:

$$\tau_2 = -\frac{T}{B}\left(\frac{B''}{2} - \frac{(B')^2}{B}\right) - \varepsilon. \qquad \text{Eq. S18}$$

Substituting $\tau_1$ and $\tau_2$ into Eq. S9 yields the general form of any (at least twice-differentiable) trade-off function for multiplicative fitness functions that satisfies the conditions of honest signalling:

$$T(q,z) = \tau_0 - \frac{\beta_1\tau_0}{\beta_0}(z-\hat{z}) - \left(\frac{\tau_0}{\beta_0}\left(\beta_2 - \frac{{\beta_1}^2}{\beta_0}\right) + \varepsilon\right)(z-\hat{z})^2 + \tau_3(z-\hat{z})^3 + \cdots. \qquad \text{Eq. S19}$$

Substituting $\tau_i$ and $\beta_j$ from Eq. S10, Eq. S16 and Eq. S18, and omitting function arguments $q$ and $z$:

$$T_M = T(q,z) = D - \frac{B'D}{B}(z-\hat{z}) - \left(\frac{D}{B}\left(\frac{B''}{2} - \frac{(B')^2}{B}\right) + \varepsilon\right)(z-\hat{z})^2 + \cdots. \qquad \text{Eq. S20}$$

Higher order terms can be anything without affecting the solution. The *equilibrium signal cost function* $D(q)$ represents the cost that signallers pay at the equilibrium. Its value can be any constant or function of $q$. Note, that in multiplicative models, where the optimum transfer of the receiver is positive ($\hat{z} > 0$), there are always two equilibrium strategy (see for example Grafen's model Fig. S4). If $D(q) > 0$, $\hat{z} > 0$ is a global optimum. If $D(q) < 0$, $\hat{z} = 0$ is the global optimum and $\hat{z} > 0$ can only be a local optimum in, coinciding with the optimum $\hat{z}$ of $w_{\mathbf{R}}$. If $D(q) = 0$, there is no single local or global optimal $z$ for any $q$, as $w_{\mathbf{S}}$ has a neutral subspace at the maximum value, spanning the region between the optimum of $w_{\mathbf{R}}$ and $z = 0$ (see Fig. S4).

Table S2 shows the first and second order Taylor coefficients for additive and multiplicative models, respectively (based on Eq. S11, Eq. S12, Eq. S16 and Eq. S18).

*Table S2. The first and second order Taylor coefficients of the trade-off function T for additive and multiplicative models, that satisfy the conditions of honest signalling. All derivatives are with respect to z, all function arguments q and z are omitted, and $\varepsilon > 0$.*

| | **First Taylor coefficient of $T$** | **Second Taylor coefficient of $T$** |
|---|---|---|
| **Additive fitness models** | $\tau_1 = -B'$ | $\tau_2 = -\frac{1}{2}B'' - \varepsilon$ |
| **Multiplicative fitness models** | $\tau_1 = -\frac{B'D}{B}$ | $\tau_2 = -\frac{D}{B}\left(\frac{B''}{2} - \frac{(B')^2}{B}\right) - \varepsilon$ |

### Appendix 4. Trade-off functions of well-known models

In this section we discuss the definitive models of the additive (Godfray 1991, Bergstrom et al. 2002) and multiplicative fitness cases (Grafen 1990), in chronological order. For a comparison of the notation and terminology of relevant models, see Table S1. For a comparison of the first and second order Taylor coefficients of these models, see Table S3.

*Table S3. The first and second order Taylor coefficients of the general trade-off function T of know models. Derivatives of B are with respect to z; $\varepsilon > 0$.*

| | **Receiver's optimal resource distribution = equilibrium transfer ($\hat{z}$)** | **Signaller's benefit function ($B$)** | **First Taylor coefficient of $T$ ($\tau_1$)** | **Second Taylor coefficient of $T$ ($\tau_2$)** |
|---|---|---|---|---|
| **Additive fitness models** | | | $\tau_1 = -B'$ | $\tau_2 = -\frac{1}{2}B'' - \varepsilon$ |
| (Godfray 1991) | $\hat{z} = \frac{\ln\left(\frac{\gamma U c}{G}\right)}{c}$ | $B = U(1 - e^{-cz}) + \psi G(Z - z)$ | $\tau_1 = cUe^{-cz} + \psi G$ | $\tau_2 = \frac{1}{2}c^2 U e^{-cz} - \varepsilon$ |
| (Bergstrom et al. 2002) | $\hat{z} = c$ | $B = z$ | $\tau_1 = -1$ | $\tau_2 = -\varepsilon$ |
| **Multiplicative fitness models** | | | $\tau_1 = -\frac{B'D}{B}$ | $\tau_2 = -\frac{D}{B}\left(\frac{B''}{2} - \frac{(B')^2}{B}\right) - \varepsilon$ |
| (Grafen 1990) | $\hat{z} = c$ | $B = c\,z^a$ | $\tau_1 = -a\,z^{-1}$ | $\tau_2 = D\left(\frac{a + a^2}{2z^2}\right) - \varepsilon$ |

#### Grafen's multiplicative model (1990)

Grafen's strategic choice multiplicative fitness model (Grafen 1990) describes a signalling game in the context of mate choice, and based on a previously proposed hypothesis for honest signalling (Zahavi 1977). Males have a hidden quality ($q$), females need to know this hidden quality to make an optimal division of resource ($z$). Males can advertise ($s$) their quality, where both the quality and the signal are continuous and differentiable. Fig. S2 shows the various functions of the model.

Receiver's (female) fitness is:

$$w_{\mathbf{R}}(q,z) = q - (z-q)^2. \qquad \text{Eq. S21}$$

The receiver's optimum transfer $z$ as a function of $c$ is (see Fig. S3):

$$\frac{\partial}{\partial z} w_{\mathbf{R}}(q,z) = 0,$$
$$\frac{\partial^2}{\partial z^2} w_{\mathbf{R}}(q,z) < 0,$$
$$\hat{z} = q.$$

Signaller's (male) fitness benefit $B$ is:

$$B(q,z) = q\, z^a. \qquad \text{Eq. S22}$$

The first and second partial derivatives of $B$ with respect to $z$ are:

$$\frac{\partial}{\partial z} B(q,z) = B'(q,z) = a\, q\, z^{a-1}, \qquad \text{Eq. S23}$$
$$\frac{\partial^2}{\partial z^2} B(q,z) = B''(q,z) = (a-1)a\, q\, z^{a-2}, \qquad \text{Eq. S24}$$

where $a = 3$ or 4 (from (Grafen 1990)).

To construct the appropriate trade-off function $T$ for $w_{\mathbf{S}} = T \cdot B$, we calculate the necessary coefficients of the Taylor series of $T$. The first Taylor coefficient is:

$$\tau_1 = -\frac{B' D(q)}{B} = -\frac{D(q)\left(a\, q\, z^{a-1}\right)}{q\, z^a} = -\frac{D(q)\, a}{z}, \qquad \text{Eq. S25}$$

where $D(q)$ can be any function of quality $c$ (or constant). There are always two equilibrium strategy, and which one is the global optimum depends on $D(q)$. If $D(q) > 0$, $z = q$ is the global optimum. If $D(q) < 0$, $z = 0$ is the global optimum and $z = c$ can only be a local optimum in the $z = q$ region (the optimum of $w_{\mathbf{R}}$). If $D(q) = 0$, there is no single local or global optimal $z$ for any $q$, as $w_{\mathbf{S}}$ has a neutral subspace at the maximum value, spanning the region between the $z = q$ optimum of $w_{\mathbf{R}}$ and $z = 0$ (see Fig. S4).

The second Taylor coefficient of $T$ is (omitting function arguments):

$$\tau_2 = -\frac{D}{B}\left(\frac{B''}{2} - \frac{(B')^2}{B}\right) - \varepsilon. \qquad \text{Eq. S26}$$

After substituting in functions of $B'$ and $B''$, the coefficient is:

$$\tau_2 = -\frac{D(q)}{q\, z^a}\left(\frac{(a-1)a\, q\, z^{a-2}}{2} - \frac{(a\, q\, z^{a-1})^2}{q\, z^a}\right) - \varepsilon =$$
$$= -D(q)\left(\frac{(a-1)a\, q\, z^{a-2}}{2(q\, z^a)} - \frac{(a\, q\, z^{a-1})^2}{(q\, z^a)^2}\right) - \varepsilon =$$
$$= -D(q)\left(\frac{(a-1)a\, z^{-2}}{2} - a^2 z^{-2}\right) - \varepsilon =$$
$$= -D(q)\left(\frac{(a-1)a - 2a^2}{2z^2}\right) - \varepsilon =$$
$$= -D(q)\left(\frac{-a - a^2}{2z^2}\right) - \varepsilon =$$

$$\tau_2 = -\left(D(q)\left(\frac{-a-a^2}{2z^2}\right) + \varepsilon\right). \quad \text{Eq. S27}$$

Substituting the Taylor coefficients Eq. S10, Eq. S25 and Eq. S27 into Eq. S9 gives the equilibrium trade-off function ($\varepsilon > 0$):

$$T(q,z) = D(q) - \frac{D(q)\,a}{\hat{z}}\ (z-\hat{z}) - \left(D(q)\left(\frac{-a-a^2}{2z^2}\right) + \varepsilon\right)\ (z-\hat{z})^2 + \cdots.$$

Equilibrium transfer is $\hat{z} = q$, thus:

$$T(q,z) = D(q) - \frac{D(q)a}{q}\ (z-\hat{z}) - \left(D(q)\left(\frac{-a-a^2}{2q^2}\right) + \varepsilon\right)(z-\hat{z})^2 + \cdots. \quad \text{Eq. S28}$$

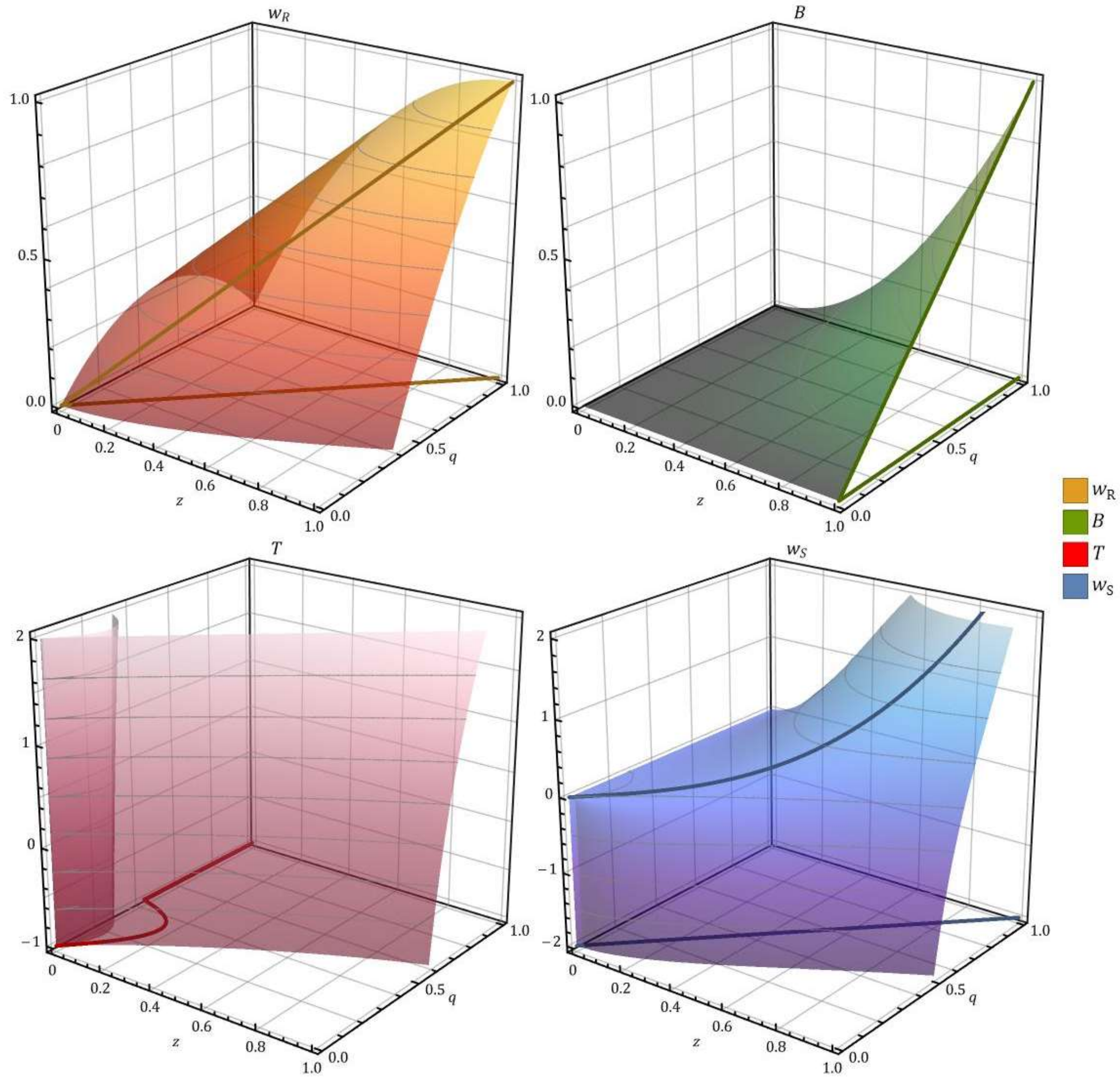


*Fig. S2. Grafen's multiplicative model. Thick lines indicate the optimal $z$ of each surface at a given $q$. Parameters are $\{a = 2, D(q) = 3, \varepsilon = 10\}$.*

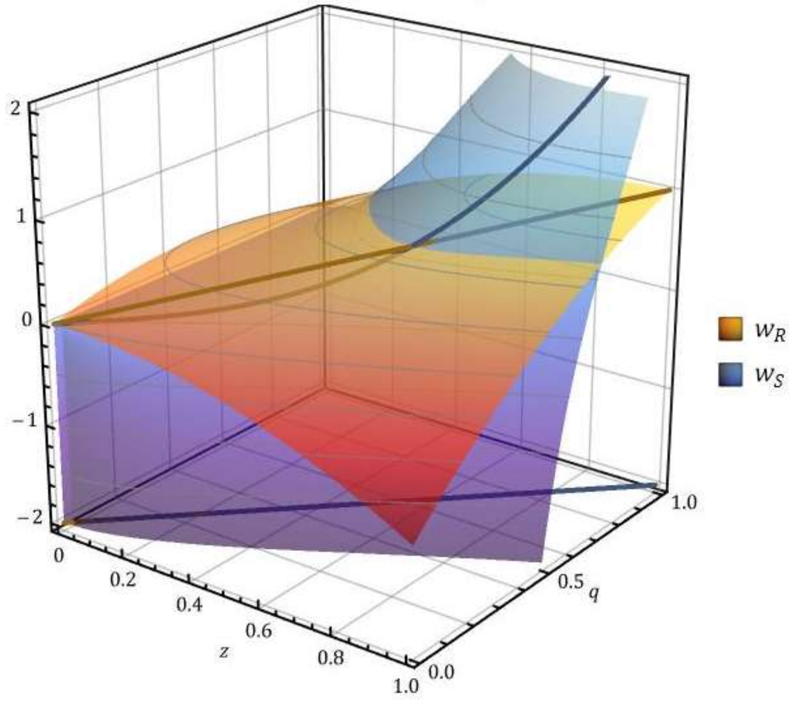


*Fig. S3. Comparison of receiver and signaller fitness functions (yellow $w_{\boldsymbol{R}}$ and blue $w_{\boldsymbol{S}}$, respectively) in Grafen's multiplicative model. Thick lines indicate the optimal $z$ of each surface at a given $q$. Notice, that the optima of the two surfaces coincide, as their projection to the $w = -2$ plane shows. Parameters are* $\{a = 2, D(q) = 3, \varepsilon = 10\}$.

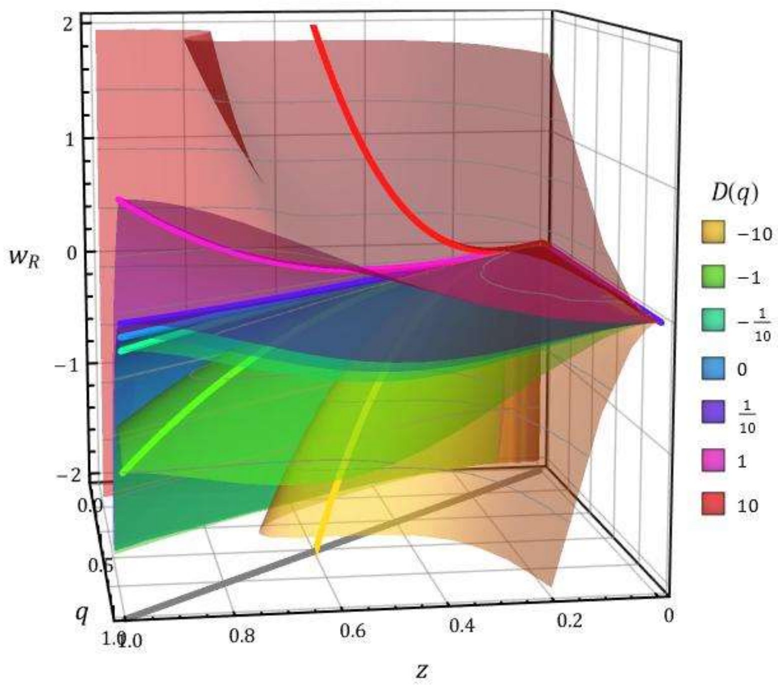


*Fig. S4. Signaller fitness function $w_{\boldsymbol{S}}$ at various constant $D(q)$ values in Grafen's multiplicative model. Thick lines indicate the optimal $z$ of each surface at a given $q$. Notice, that the optima of all surfaces coincide with the optima of $w_{\boldsymbol{R}}$ in the $(q, z)$ plane, indicated as a grey line projected to the plane at $w_{\boldsymbol{R}} = -2$. Parameters are* $\{a = 2, \varepsilon = 10\}$.

#### Godfray's additive model (1991)

Godfray's model (Godfray 1991) differs from Grafen's model in three respects. First, it considers a differential benefit instead of a differential cost model, i.e., the cost of signalling as a function of signal intensity ($x$) is the same for all signallers; however, the value of benefit is conditional on the quality of the signaller ($q$, originally condition $c$ in (Godfray 1991)). Second, Godfray investigated a model parent-offspring communication, i.e., a context where there is a relatedness between signaller and receiver. This relatedness changes the benefit function for both signaller and receiver. Third, fitness components are additive. Fig. S5 shows the various functions of the model.

The fitness functions of signaller and receiver can be written as follow:

$$w_{\mathbf{S}}(q,x,z) = h(q,z) - f(x) + \psi g(Z - z),$$
$$w_{\mathbf{R}}(q,x,z) = \gamma(h(q,z) - f(x)) + g(Z - z),$$

where the first term gives the direct fitness benefit, the second term gives the cost of signals, and the third term gives the benefit due to relatedness.

$$h(q,z) = U(1 - e^{-qz}),$$
$$g(Z - z) = G(Z - z).$$

Receiver's (female's) fitness is:

$$w_{\mathbf{R}}(q,z) = \gamma U(1 - e^{-qz}) + G(Z - z).$$

The receiver's optimum transfer $z$ as a function of $q$ is (see Fig. S6):

$$\hat{z} = \frac{\ln\left(\frac{q\gamma U}{G}\right)}{q}.$$

Signaller's (offspring's) fitness benefit $B$ is:

$$B(q,z) = U(1 - e^{-q} \ ) + \psi G(Z - z).$$

The first and second partial derivatives of the signaller's benefit function, respectively:

$$\frac{\partial}{\partial z}B(q,z) = -qU\,e^{-q} \ - \psi G,$$
$$\frac{\partial^2}{\partial z^2}B(q,z) = -q^2U\,e^{-qz}.$$

Accordingly, the first and second Taylor coefficients are:

$$\tau_1 = -\,B'(\hat{z}) = qUe^{-qz} + \psi G,$$
$$\tau_2 = -\frac{1}{2}B''(\hat{z}) - \varepsilon = \frac{1}{2}q^2U\,e^{-qz} - \varepsilon,$$

where $\varepsilon > 0$. Substituting the above $\tau_i$ into Eq. S9, the equilibrium trade-off function is:

$$T(q,z) = D(q) + (qUe^{-qz} + \psi G)\,(z - \hat{z}) + \left(\frac{1}{2}q^2Ue^{-qz} - \varepsilon\right)\,(z - \hat{z})^2 + \cdots.$$

Equilibrium transfer is $\hat{z} = \ln\left(\frac{\gamma Uq}{G}\right)/q$ , thus:

$$T(q,z) = D(q) + \left(qU\exp\left(-\frac{q\ln\left(\frac{\gamma Uq}{G}\right)}{q}\right) + \psi G\right)\left(z - \frac{\ln\left(\frac{\gamma Uq}{G}\right)}{q}\right) +$$
$$+\left(\frac{1}{2}q^2U\exp\left(-\frac{q\ln\left(\frac{\gamma Uq}{G}\right)}{q}\right) - \varepsilon\right)\left(z - \frac{\ln\left(\frac{\gamma Uq}{G}\right)}{q}\right)^2 + \cdots,$$

where $\varepsilon > 0$. In Godfray's cost-free model, $D(q) = 0$.

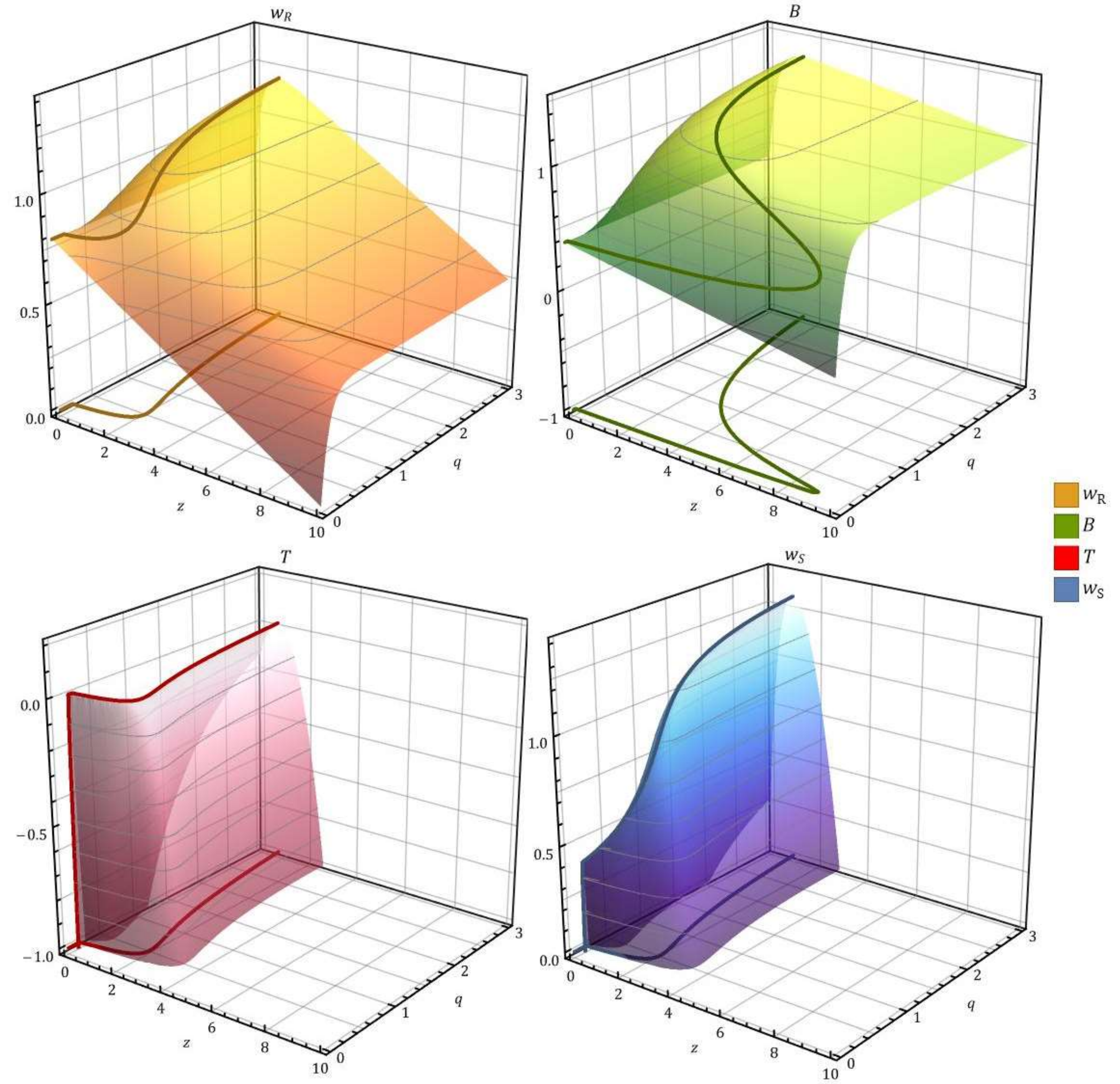


*Fig. S5. Godfray's additive model* (Godfray 1991)*. Thick lines indicate the optimal $z$ of each surface at a given $q$. Parameters are* $\{\psi = 1/2, \gamma = 1/2, G = 0.08, U = 1, Z = 10, D(q) = 0, \varepsilon = 1\}$.

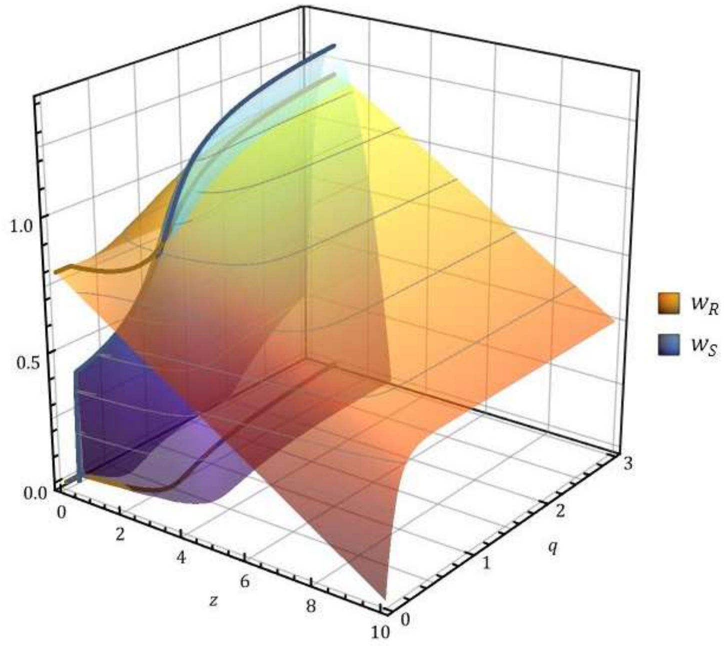


*Fig. S6. Comparison of receiver and signaller fitness functions (yellow $w_{\mathbf{R}}$ and blue $w_{\mathbf{S}}$, respectively) in Godfray's additive model* (Godfray 1991). *Thick lines indicate the optimal $z$ of each surface at a given $q$. Notice, that the optima of the two surfaces coincide, as their projection to the $w = -2$ plane shows. Parameters are* $\{\psi = 1/2\,, \gamma = 1/2\,, G = 0.08, U = 1, Z = 10, D(q) = 0, \varepsilon = 1\}$.

#### Bergstrom et al.'s additive model (2002)

At the additive model of (Bergstrom et al. 2002), receiver's (female) fitness is (see Fig. S7):

$$w_{\mathbf{R}}(q,z) = e^{-(q-z)^2}.$$

The receiver's optimum transfer $z$ as a function of $q$ is (see Fig. S8):

$$\hat{z} = q.$$

Signaller's (male) fitness benefit $B$ is:

$$B(q,z) = z.$$

The first and second partial derivatives of the signaller's benefit function, respectively:

$$\frac{\partial}{\partial z}B(q,z) = 1,$$
$$\frac{\partial^2}{\partial z^2}B(q,z) = 0.$$

Accordingly, the first and second Taylor coefficients at $z = \hat{z}$ are:

$$\tau_1 = -\,B' = -1,$$
$$\tau_2 = -\frac{1}{2}B'' - \varepsilon = -\varepsilon.$$

Substituting $\tau_0, \tau_1, \tau_2$ into Eq. S9, the equilibrium cost is:

$$T(q,z) = D(q) + 1(z - \hat{z}) + (-\varepsilon)(z - \hat{z})^2 + \cdots.$$

Equilibrium transfer is $\hat{z} = q$, thus:

$$T(q,z) = D(q) + (z - q) - \varepsilon(z - q)^2 + \cdots,$$

where $\varepsilon > 0$.

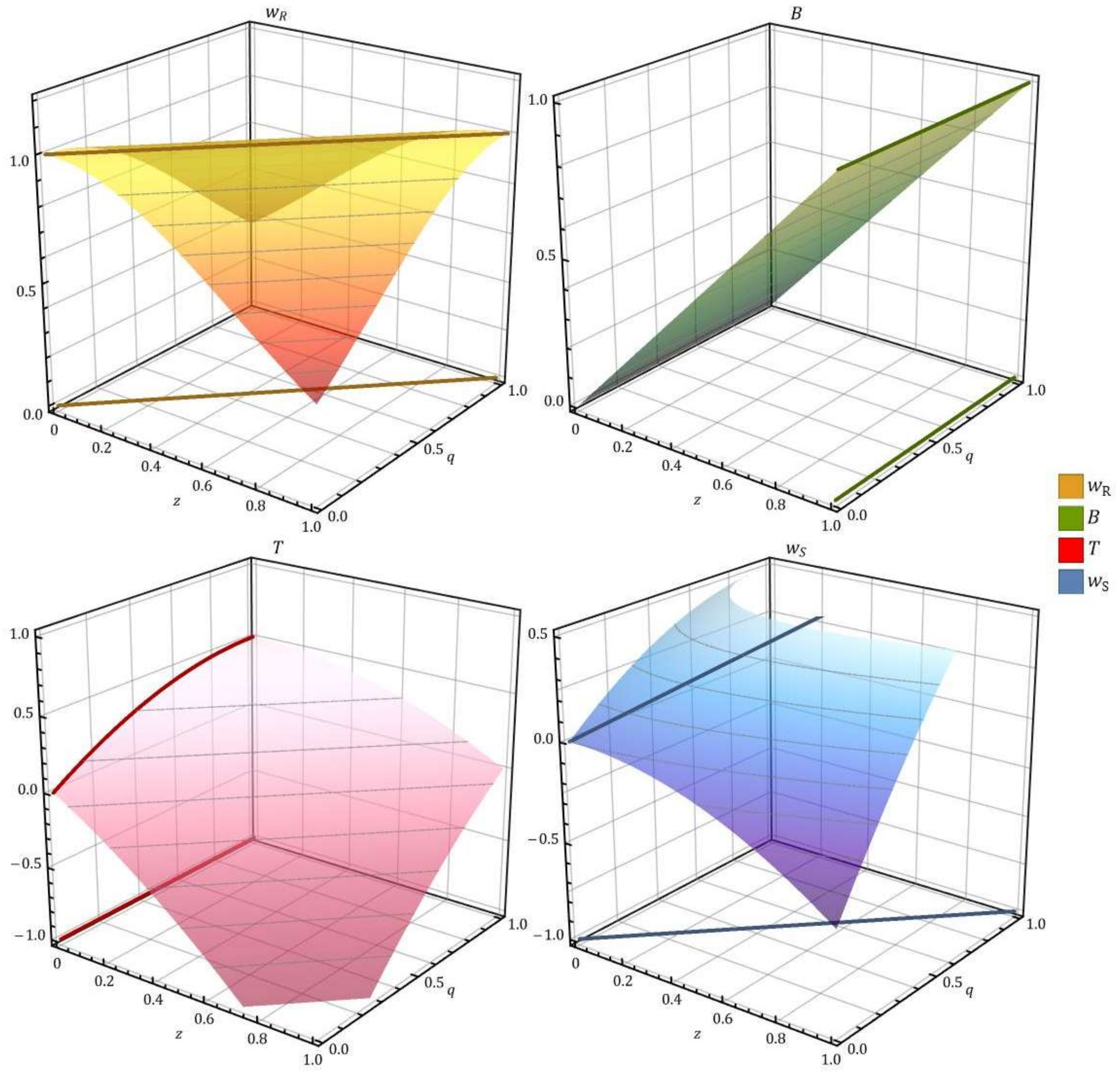


*Fig. S7. Bergstrom et al.'s additive model* (Bergstrom et al. 2002). *Thick lines indicate the optimal $z$ of each surface at a given $q$. Parameters are $\{D(q) = 0, \varepsilon = 1\}$.*

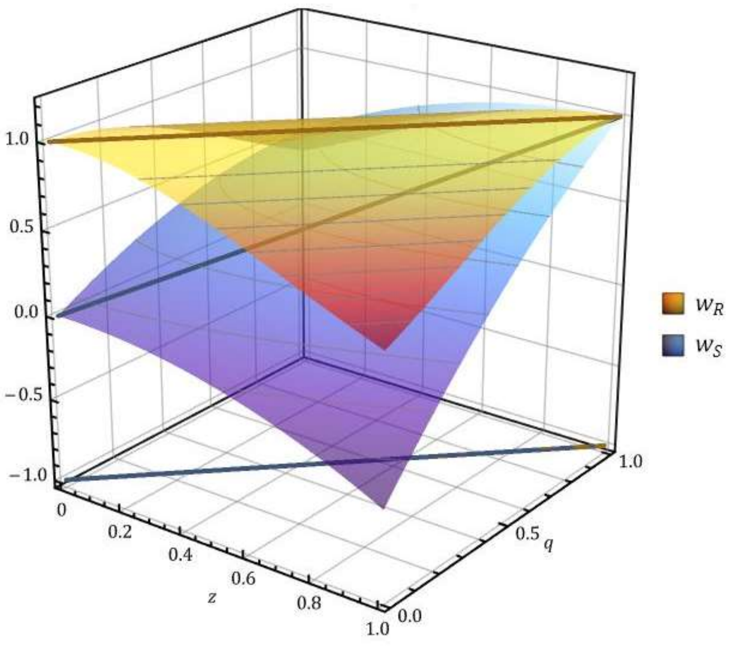


*Fig. S8. Comparison of receiver and signaller fitness functions (yellow $w_{\boldsymbol{R}}$ and blue $w_{\boldsymbol{S}}$, respectively) in Bergstrom et al.'s additive model* (Bergstrom et al. 2002). *Thick lines indicate the optimal $z$ of each surface at a given $q$. Notice, that the optima of the two surfaces coincide, as their projection to the $w = -2$ plane shows. Parameters are $\{D(q) = 0, \varepsilon = 1\}$.*

#### Appendix 5. Single step optimization models of signalling

In many models signalling is investigated as a single step optimization problem from the point of view of the signaller while the receiver's strategy is assumed to be fixed (Holman 2012, Biernaskie et al. 2014, Clifton et al. 2016, Fromhage and Henshaw 2022). While this approach can greatly reduce the technical complexity of the solution there is no guarantee that the assumed receiver response is an optimal response (best strategy) against the calculated signaller strategy. As a result, there is no guarantee that the identified signaller-receiver strategy pair is evolutionarily stable. A further problem with these models is that the receiver's fitness is not even defined (thus the equilibrium path cannot be calculated), which would be necessary to solve the double optimization problem posed by signalling. Therefore, conclusions drawn from these models should be considered only as preliminary results, interpreted with caution.

Holman has investigated the strategic cost of pheromone signalling in ants (Holman 2012). Unfortunately, his model is not a complete signalling game as he assumed a fixed receiver response, independent of the signal. A unifying model of costly signalling was proposed by Biernaskie and colleagues (Biernaskie et al. 2014). They were able to show that the previous distinction between so called "indices" and "costly signals" (aka. handicaps) is quantitative and not qualitative. Unfortunately, their model is based on the model of (Holman 2012), which is not a signalling game. This, in turn implies that the variant of the model investigated by (Biernaskie et al. 2014) is not a signalling game either for the same reason explained above. Clifton and colleagues have investigated the evolution of dimorphic features (features with bimodal optima) (Clifton et al. 2016). They assumed a pre-set trade-off function and they further assumed a 'playing the field'-type fitness function, i.e. where the fitness of the ego depends on the average type found in the population. This latter assumption is crucial to achieve the bimodal result (see (Számadó and Penn 2018)). Unfortunately, they assumed a fixed response for the receiver, and they did not investigate whether the receiver's fixed strategy is evolutionarily stable (is best response) against the identified bimodal signaller strategy.

#### Appendix 6. The model of (Biernaskie et al. 2018)

Biernaskie and colleagues have investigated a variant of Grafen's seminal model (Grafen 1990) to investigate the effect of signal evolution on signal cost (Biernaskie et al. 2018). They used a multiplicative fitness function with a pre-defined trade-off function and exponential benefits. Note that both the receiver's fitness function ($w_{\mathbf{R}}$) and the signaller's benefit function ($B$) are the same as in Grafen's original model, thus the same general solution applies to this model as well (Eq. S28). Biernaskie and colleagues picked a specific trade-off function to serve as an example. While their solution is formally correct, it does not represent a general solution to their model.

#### Appendix 7. The model of (Fromhage and Henshaw 2022)

Fromhage and Henshaw (Fromhage and Henshaw 2022) has investigated the first condition of our model (Eqs. S11, S16) but 1) without the context of the signalling game and 2) without the Taylor series decomposition to reverse-engineer a general solution. Point 1) implies that they did not evaluate the first condition (extremum condition, Eq. S3) at the optimum of the receiver, and they did not evaluate the second condition (stability condition, Eq. S4) at all. They conclude that multiplicative fitness functions are more likely to promote honesty (as opposed to additive ones). Unfortunately, due to the above-mentioned shortcomings, their result is misleading. Our results show, that both additive and multiplicative fitness functions can maintain honest signalling.

#### Appendix 8. The 'Lazy Student' game

Students are preparing for an exam. They have two time slots for preparing, with the second slot coinciding with a concert. The exam is very important, and students cannot afford to fail it. On the other hand, the concert is free and enjoyable (compared to any exam). There are two types of students: Prepared students efficiently prepare for the exam using the first time slot only. Lazy students do nothing during the first time slot. The question is: under what condition is attending the concert (and brandishing this fact as a signal) conveys information about the type of the student? See Fig. S9 for a visual description o the game.

It is clear that Prepared students can attend the concert without jeopardizing their exam, as they have already prepared for it. Lazy students, however, must make a strategic decision, having only one free timeslot left. They must choose between attending the concert or preparing for the exam. If Lazy students cannot afford to fail the exam (too much cost), they have to trade in the concert for preparing for the exam. In this scenario, visiting the concert conveys information about the type of the student.

Note, that at the honest equilibrium, no one pays the cost of failing the exam. High quality

(Prepared) individuals do not waste resources for costly ('handicap') signals, in fact they can enjoy the free concert, which therefore effectively serves as a beneficial signal. If anything is wasted in this game at the equilibrium, it is the opportunity of the Lazy student to visit a free concert. Thus, contrary to Zahavi's prediction, a beneficial signal can be honest and stable at equilibrium. Prepared (honest) students are efficient while Lazy students (potential cheaters) are inefficient (being wasteful).

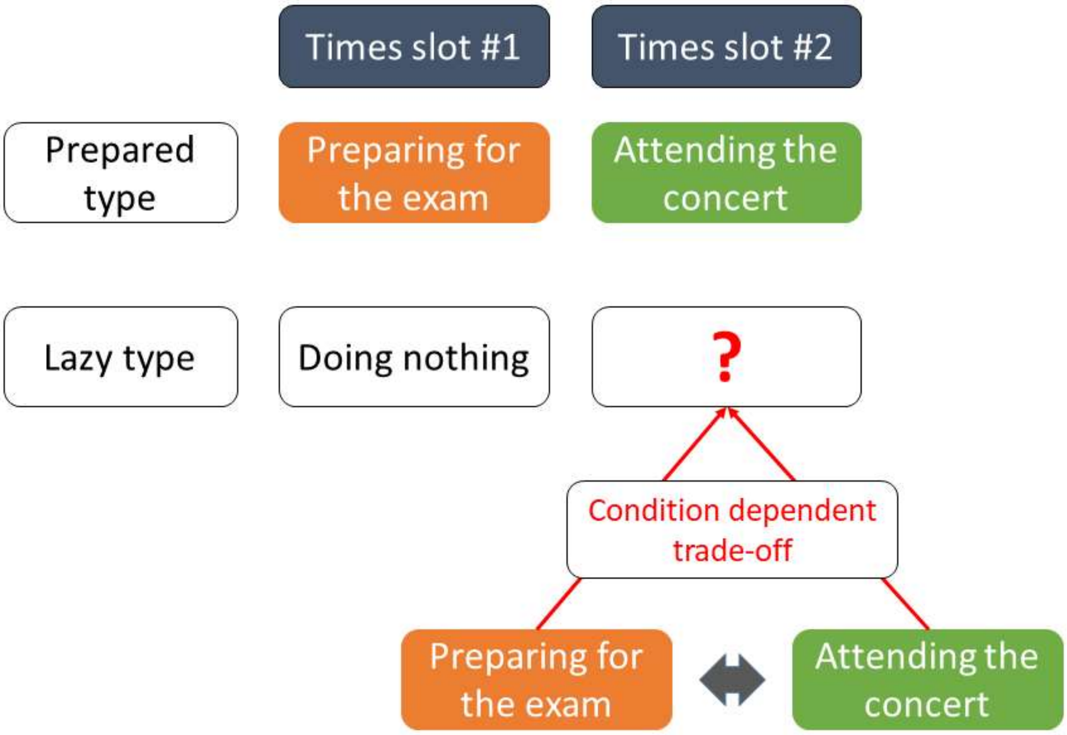


*Fig. S9. The 'Lazy Student' game. There are two time slots for the students to use, three activities ('doing nothing', 'preparing for the exam', 'attending the concert') and two types of students. 'Prepared' students use their time slots efficiently therefore they can attend the concert. 'Lazy' students waste the first time slot by doing nothing. As a result, they must face a trade-off between 'preparing for the exam' or 'attending the concert'. Concert attendance is an honest signal of student type if the marginal cost of attending the concert for 'Lazy' students (failing the exam) is higher than the marginal benefit (enjoying the concert).*
